# Converting an allocentric goal into an egocentric steering signal

**DOI:** 10.1101/2022.11.10.516026

**Authors:** Peter Mussells Pires, L.F. Abbott, Gaby Maimon

## Abstract

Neuronal signals relevant for spatial navigation have been described in many species^1–12^, however, a circuit-level understanding of how such signals interact to guide behaviour is lacking. Here we characterize a neuronal circuit in the *Drosophila* central complex that compares internally generated estimates of the fly’s heading and goal angles––both encoded in world-centred, or allocentric, coordinates––to generate a body-centred, or egocentric, steering signal. Past work has argued that the activity of EPG cells, or “compass neurons”^2^, represents the fly’s moment-to-moment angular orientation, or *heading angle*, during navigation^13^. An animal’s moment-to-moment heading angle, however, is not always aligned with its *goal angle*, i.e., the allocentric direction in which it wishes to progress forward. We describe a second set of neurons in the *Drosophila* brain, FC2 cells^14^, with activity that correlates with the fly’s goal angle. Furthermore, focal optogenetic activation of FC2 neurons induces flies to orient along experimenter-defined directions as they walk forward. EPG and FC2 cells connect monosynaptically to a third neuronal class, PFL3 cells^14,15^. We found that individual PFL3 cells show conjunctive, spike-rate tuning to both heading and goal angles during goal-directed navigation. Informed by the anatomy and physiology of these three cell classes, we develop a formal model for how this circuit can compare allocentric heading- and goal-angles to build an egocentric steering signal in the PFL3 output terminals. Quantitative analyses and optogenetic manipulations of PFL3 activity support the model. The biological circuit described here reveals how two, population-level, allocentric signals are compared in the brain to produce an egocentric output signal appropriate for the motor system.

## Introduction

Dung beetles pick an arbitrary direction in which to roll their precious ball of dung^16^. Fruit bats fly kilometers to re-visit the same tree night after night^17^. Whether their goal is to reach a specific location in space, like bats, or to maintain a consistent angular bearing, like dung beetles, animals must regularly update their locomotor behaviour (e.g. turn left or right) based on whether they are heading in the correct direction.

To determine which way to turn during navigation, the brain could compare an explicit internal estimate of the animal’s *heading angle* (i.e., the animal’s moment-to-moment orientation, or compass direction) with a *goal angle*^13,18^ (i.e., the compass direction along which an animal wishes to progress forward). The difference between these two angles could then direct turns toward the goal (Fig. 1a). Heading and goal angles are closely related because animals typically orient in the direction in which they wish to progress forward; however, the two angles are distinct because the goal angle remains constant in the face of occasional turns or detours that briefly change the animal’s heading angle. Importantly, when heading and goal angles are both encoded in a common, allocentric or world-referenced (e.g., north/east/ south/west) coordinate frame, a neural circuit that compares them appropriately would yield a signal in egocentric or body-referenced (e.g., left/right) coordinates appropriate for determining the direction and vigor of steering.

**Fig. 1.**
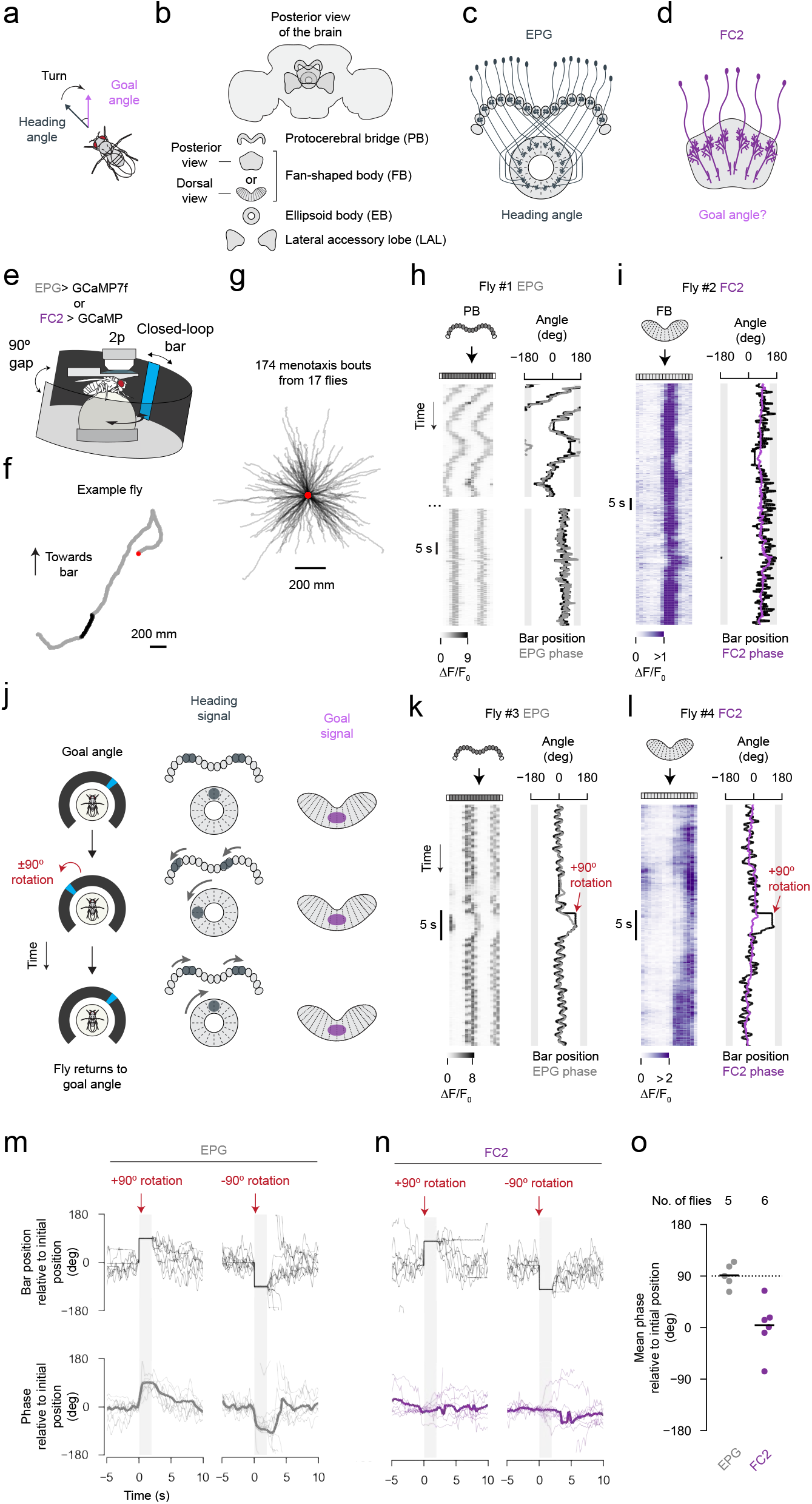
FC2 neurons express a stable activity bump in the fan-shaped body during virtual rotations of the fly. **a**, Comparing heading and goal angles to drive turning. **b**, Schematic of three central-complex structures and the lateral accessory lobes of the fly brain. **c**, Schematic of EPG neurons. **d**, Schematic of FC2 neurons. **e**, Virtual reality setup for recording neural activity in a walking fly. **f**, Virtual 2D trajectory from a single fly performing menotaxis from whom we simultaneously recorded GCaMP activity (26 min. recording). Note that for some portions of the trajectory, the fly traveled in the same direction for hundreds of body lengths. We algorithmically extracted relatively straight segments of the fly’s trajectory—which we refer to as menotaxis bouts—one of which is highlighted in black (see Methods). Red dot marks the start of the trajectory. **g**, Trajectories of all menotaxis bouts from the FC2 and EPG imaging data set analyzed in this figure. Trajectories were aligned to begin at the same location (red dot). **h**, Example trace of jGCaMP7f activity of EPG neurons in the protocerebral bridge. Left: EPG ΔF/F0 over time. Right: Bar position (i.e. the inverse of the fly’s heading angle) (black) and the EPG phase estimate (grey). Shaded area represents the 90° gap where the bar is not visible. The top trace shows a time period where fly meandered rather than performing menotaxis. The bottom trace shows a later moment, when the same fly maintained a relatively consistent heading angle while walking forward (forward velocity not shown). Note that the EPG phase tracks the bar position throughout, consistent with previous studies^2,13^. **i**, Example trace of jGCaMP7f activity of FC2 neurons in the fan-shaped body (viewed dorsally) from a fly performing menotaxis (i.e. stabilizing a relatively consistent heading as it walks forward). **j**, Left: Experimental paradigm for dissociating heading and goal signals (see text). **k**, Example EPG trace during a +90° virtual rotation. The EPG bump tracks the virtual rotation. **l**, Example FC2 trace during a +90° virtual rotation. The FC2 bump does not track the virtual rotation. **m**, Individual ±90° rotation trials. Top: bar position zeroed at onset of rotation. Bottom: EPG phase zeroed at onset of rotation; thick lines show the mean across flies. Only trials where flies turned so as to realign their heading with the heading angle prior to the bar jump were analyzed, because it is in these trials that we could be most confident that the fly’s goal did not change during the bar jump (see Methods for trial selection criteria). Seventeen ±90° trials from 5 flies are shown. Shaded area marks the 2 s period when the bar was kept stable, at a ±90° offset, before giving the fly closed-loop control. **n**, Same as panel **m** but for fifteen ±90° trials from 6 FC2 flies. **o**, Mean phase value during final 1 s of the open-loop period in panels **m** and **n**. Each dot is the mean for one fly. Horizontal lines show the mean across flies. The sign of the phase was flipped for −90° trials in order to combine +90° and −90° trials together for each fly. Dashed line shows the expected phase position after the bar jump if a bump’s position in the brain were to track the bar angle. V-test for FC2 flies: μ=0°, p=2.58×10-3. V-test for EPG flies: μ=90°, p=2.62×10-4.

Neural signals relevant for such a computation have been described in many species. For example, neural correlates of moment-to-moment heading (i.e. head-direction cells) exist in vertebrates^1,19,20^ and invertebrates^2,21,22^ as do neurons with activity related to navigational goals^10,11,23,24^ and locomotor turns^25–27^. Yet, despite these correlates and associated computational models for goal-directed navigation^18,27–30^ a biological circuit that converts allocentric, navigation-related signals into an output appropriate for the motor system has yet to be described.

We probed the neurophysiology of a navigational circuit in fruit flies maintaining a persistent angular bearing^13,31^. Whereas a previously described neural population tracks the flies’ heading angle during this simple task^13^, we describe a second neural population whose activity correlates with, and can determine, the flies’ goal angle. Via patch-clamp electrophysiology, two-photon imaging, computational modeling and neuronal perturbations, we show how a neuronal cell type, monosynaptically downstream of the above two cell types, compares allocentric heading- and goal-angle inputs to produce an egocentric steering-signal. This circuit allows a fly to navigate in the world along any desired compass direction.

### Central complex and menotaxis

The insect central complex is a set of midline-straddling brain structures that include the ellipsoid body, protocerebral bridge, and fan-shaped body^32^ (Fig. 1b). *Columnar neurons* of the central complex innervate subsections or columns of larger structures, with each columnar cell class tiling the structure(s) they innervate^14,33–35^. EPG cells are a class of columnar neurons that tile the ellipsoid body with their dendrites and the protocerebral bridge with their axons^35^ (Fig. 1c). EPG cells have been referred to as “compass” neurons because they express a bump of calcium activity in the circular ellipsoid body, and two copies of that bump in the linear protocerebral bridge, with the position of these bumps in the bridge or ellipsoid body (i.e., their *phase*) tracking the fly’s allocentric heading angle^2,13^. Might there exist an allocentric goal angle signal in the central complex that could be compared with the EPG heading signal to guide navigation? Inspired by past theoretical work^18,27,29^, we hypothesized that columnar neurons of the fan-shaped body might signal the fly’s goal angle. Specifically, we found that FC2 cells––a class of columnar neurons that receive inputs and send outputs within the fan-shaped body^14,15^ (Fig. 1d)––could serve such a role.

We performed two-photon calcium imaging in tethered flies, while they walked on an air-cushioned ball in a simple virtual environment^36–38^ (Fig. 1e) (Methods). The environment consisted of a vertical blue bar displayed on a panoramic LED display^39^. The bar rotated in angular closed-loop with the fly’s yaw rotations (i.e. left/right turns), thus simulating a fixed, distant cue, like the sun, whose position on the arena could be used by the fly to infer its heading in the virtual world. In this setup, we have found that flies can be motivated to walk forward for many hundreds of body lengths along a stable but seemingly arbitrary bearing relative to the visual cue^13^ –– a behaviour called *arbitrary-angle fixation* or *menotaxis*^13,31,40^. Previous work showed that menotaxis is an EPG-dependent behaviour^13,31^ and that the EPG phase encodes the fly’s *heading angle* during this task^13^.

### FC2 cells signal a goal angle

We imaged GCaMP7^41^ fluorescence from EPG and FC2 neurons (Extended Data Fig. 1) as flies performed menotaxis. We focused on time periods in which flies were stabilizing a consistent angle while walking forward (Fig. 1f, black highlight in trajectory, Fig. 1g, Extended Data Fig. 2) (Methods). During such *menotaxis bouts*, we could be confident that the flies were in a consistent behavioural state.

Much as EPG cells express bumps of activity that shift around the ellipsoid body and protocerebral bridge^2,38,42^ (Fig. 1h top), we found that FC2 cells express a calcium bump that shifts across the left/right axis of the fan-shaped body (Fig. 1i and Extended Data Fig. 3a-c). Both the EPG and the FC2 bumps had a phase that generally correlated with the position of the bar over the course of a recording (EPG: r=0.60, FC2: r=0.42,), which would be expected for bumps that track either the heading or goal angles. During menotaxis bouts, when flies were stabilizing a specific heading angle, we observed that both the EPG and FC2 bumps remained at a relatively stable position (Fig. 1h bottom & Fig. 1i). To dissociate whether the FC2 and EPG bumps better track the goal or heading angle, we virtually rotated flies ±90° while they performed menotaxis. Specifically, we discontinuously jumped the bar, in open-loop, and then returned the system to closed-loop control after a two second delay. Following such rotations, flies typically (but not always) slow their forward velocity and make a corrective turn to realign themselves with their previous heading angle^13^ (Extended Data Fig. 4). We reasoned that the fly’s goal had stayed constant throughout this perturbation on trials where the fly clearly returned to its previous heading (Methods). On such trials, heading and goal signals are expected to behave differently: a bump that tracks the heading angle should rotate ±90° and a bump that tracks the goal angle should remain fixed (Fig. 1j). We found that the EPG phase, on average, rotated approximately ±90°, in lock step with the fly’s heading, during virtual rotations of the fly, whereas the FC2 phase, on average, did not measurably deviate (Fig. 1j-o). These results are consistent with the EPG phase signaling the allocentric heading angle and the FC2 phase signaling the allocentric goal. We observe similar results independently of how we select trials for analysis (Extended Data Fig. 3d-f). If the FC2 bump can indeed signal the fly’s goal angle to downstream circuits, experimentally repositioning the FC2 bump to different left/right positions along the fan-shaped body should induce flies to walk along experimenter-defined goal directions. We next tested this hypothesis.

### Experimentally controlling the goal angle

We optogenetically activated FC2 neurons in a contiguous subset of fan-shaped body columns while monitoring the fly’s walking behaviour (Fig. 2a, Extended Data Fig. 5). Specifically, we co-expressed the red-shifted channelrhodopsin CsChrimson^43^ and sytGCaMP7f^4^ in FC2 neurons and used a two-photon laser to repeatedly reposition the FC2 bump at one of two locations, separated by approximately half the width of the left/right axis of the fan-shaped body (Fig. 2b, Extended Data Fig. 5b). If the position of the FC2 bump in the fan-shaped body signals the fly’s goal direction, this perturbation should cause a fly to repeatedly switch its heading between two angles separated by ∼180° (Extended Data Fig. 5e,f). Remarkably, flies tended to stabilize a consistent heading angle when we stimulated a given region of the fan-shaped body (Fig. 2c,e, and Extended Data Fig. 5g). Moreover, the behavioural angles flies stabilized for the two stimulation locations differed by ∼173°, on average, similar to the ∼180° predicted from the anatomical stimulation locations (Fig. 2c,e-g, Extended Data Fig. 5e). Control flies that did not express CsChrimson showed no measurable change in FC2 calcium activity during stimulations (Extended Data Fig. 5c) and did not reliably adopt the same heading direction for consistent stimulation locations (Fig. 2d-e). The behavioural heading distributions for control flies showed more overlap between the two stimulation locations (Fig. 2e-f), as expected from the fact that flies are unlikely to spontaneously flip-flop between two goal angles, 180° apart.

**Fig. 2.**
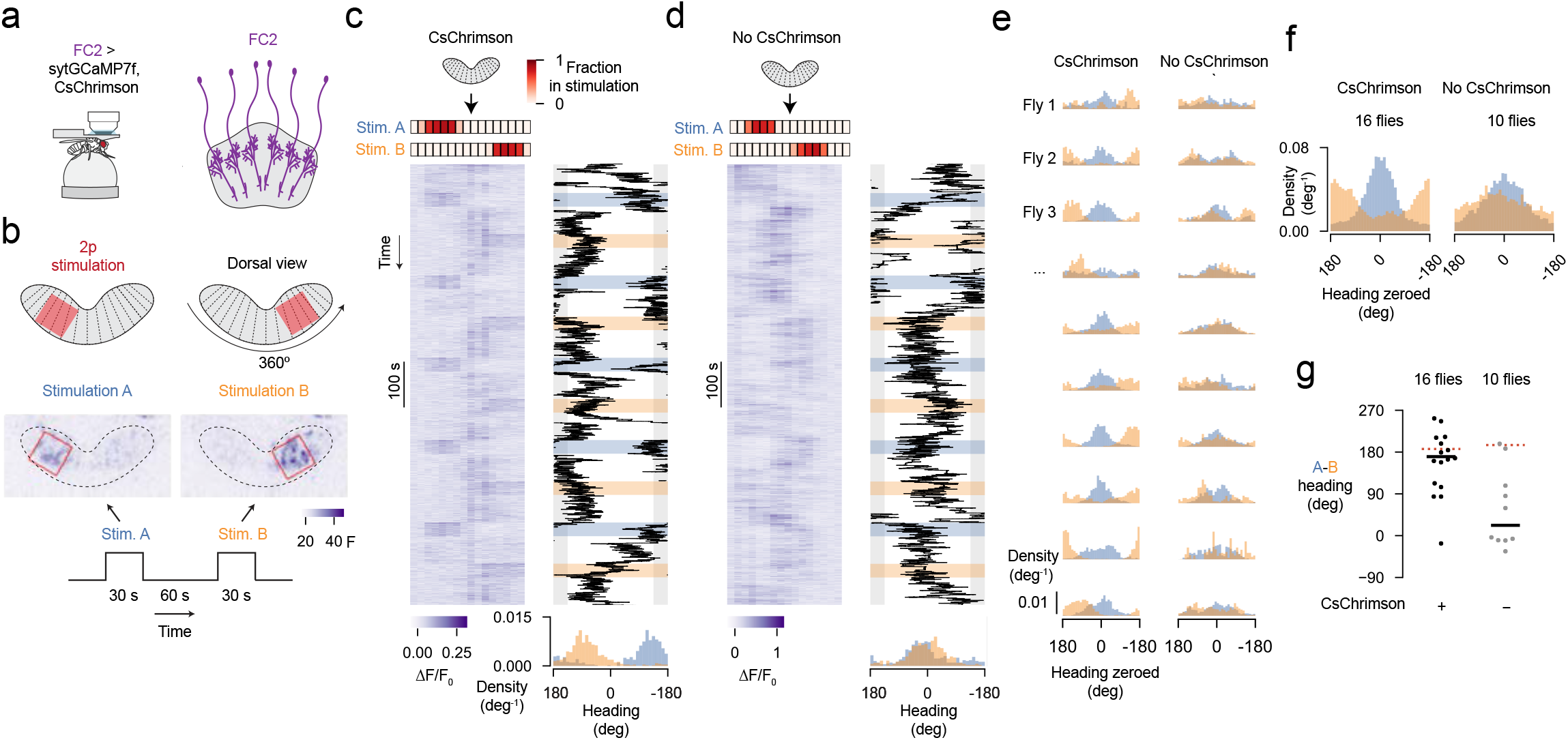
Stimulating FC2 neurons in a contiguous subset of fan-shaped body columns induces flies to orient along defined goal angles. **a**, Simultaneous imaging and focal stimulation of FC2 neurons. **b**, Stimulation protocol. Images show average *z*-projection of raw fluorescence signal during the stimulation period from a single trial. Red squares mark the two stimulation locations in the fan-shaped body (referred to as stim. A and stim. B). **c**, Example FC2 ΔF/F0 signal and behavioral traces during a CsChrimson experiment. Left: FC2 activity over time. Red heatmap shows the fraction of pixels of each column’s region of interest (ROI) that are inside the stimulation ROI. Right: fly’s heading direction over time. Shaded blue and orange areas indicate the stimulation period. Bottom: probability distribution of the fly’s heading direction across all trials for each stimulation location. **d**, Same as panel **c** but for a control fly that did not express CsChrimson. **e**, Probability distributions of heading direction for 10 (of the 16 total) CsChrimson-expressing flies (left) and 10 (of 10) control flies, which did not express CsChrimson (right). The heading direction was zeroed by subtracting the fly’s mean heading direction across all “stim. A” trials. **f**, Mean probability distributions for all flies. **g**, Difference between mean heading direction during stim. A and stim. B trials for each fly (black dots). Horizontal black line indicates the mean across flies. Dashed red line indicates the expected difference in heading direction based on the mean difference in the stimulation location for each group (see Extended Data Fig.9e). V-test for CsChrimson flies: μ=-173.4° (left dashed line), p=1.15×10-3. V-test for no CsChrimon flies: μ=-164.8° (right dashed line), p=0.932.

Previous work has shown that each fly learns an idiosyncratic offset between its heading (relative to the bar position) and its EPG phase^2^ such that for one fly the EPG bump might be at the top of the ellipsoid body when the bar is directly in front and for another fly the bump might be at the bottom. Likewise, for a given FC2 stimulation location in the fan-shaped body, individual experimental flies stabilized a consistent goal angle relative to the bar, but the value of this angle differed from fly to fly (Extended Data Fig. 5g-h). Because past work has shown that the fan-shaped body inherits its azimuthal reference frame from EPG cells^4^, these data are consistent with the FC2 phase encoding the fly’s goal angle in the same, allocentric reference frame used by the EPG neurons to encode the fly’s heading. Overall, these results provide further evidence that FC2 neurons can communicate a goal angle, in allocentric coordinates, to downstream neurons to guide behaviour.

### Feedback inhibition in FC2 cells

Stimulation of FC2 neurons in specific columns of the fan-shaped body was accompanied by a decrease of calcium signal in non-stimulated columns (Extended Data Fig. 5c). The further away an FC2 column was from the stimulation site, the larger was its decrease in activity (Extended Data Fig. 5d). This result suggests that active FC2 cells inhibit less active FC2 cells, perhaps for the purpose of promoting a single bump of activity, or a single goal angle, in their population activity pattern at any one time.

### Conjunctive tuning to heading and goal-angles in PFL3 cells

Given that EPG and FC2 cells have activity associated with the fly’s heading and goal angles, respectively, how might these two signals be compared to guide locomotion? It has been suggested that PFL3 cells^14,27,44^, a columnar cell class with compelling anatomy, could perform a heading-to-goal comparison^14,18,27,45^.

PFL3 cells receive input synapses in the protocerebral bridge and fan-shaped body, and express output synapses in the lateral accessory lobes (LALs)^14,15^, which symmetrically flank the central complex (Fig. 3a). In the bridge, PFL3 cells receive extensive synaptic input from EPG cells^14,15^, where they can thus receive signals related to the fly’s heading angle (Extended Data Fig. 6a,b,d). PFL3 neurons also receive disynaptic EPG input in the bridge, via a set of local interneurons called Δ7 cells^14,15^ (Extended Data Fig. Fig. 6c,e); Δ7 input could alter, in subtle but functionally important ways, the heading-tuning of PFL3 neurons^4^. PFL3 cells also receive strong synaptic input from FC2 neurons in the fan-shaped body^14,15^ (Extended Data Fig. 7) and thus they could receive goal-angle related information there. Individual PFL3 neurons project to either the left or right LAL where they synapse onto descending neurons (i.e., neurons connecting the brain to the ventral nerve cord) involved in steering behaviour^14,15,27^ (Fig. 3a). We will define ‘left’ and ‘right’ PFL3 neurons based on the side of the LAL to which a given neuron projects (which is typically, but not always, opposite to the side of their innervation in the protocerebral bridge). PFL3 neurons thus seem perfectly poised to compare heading inputs in the bridge with goal inputs in the fan-shaped body to impact steering signals in the LAL.

**Fig. 3.**
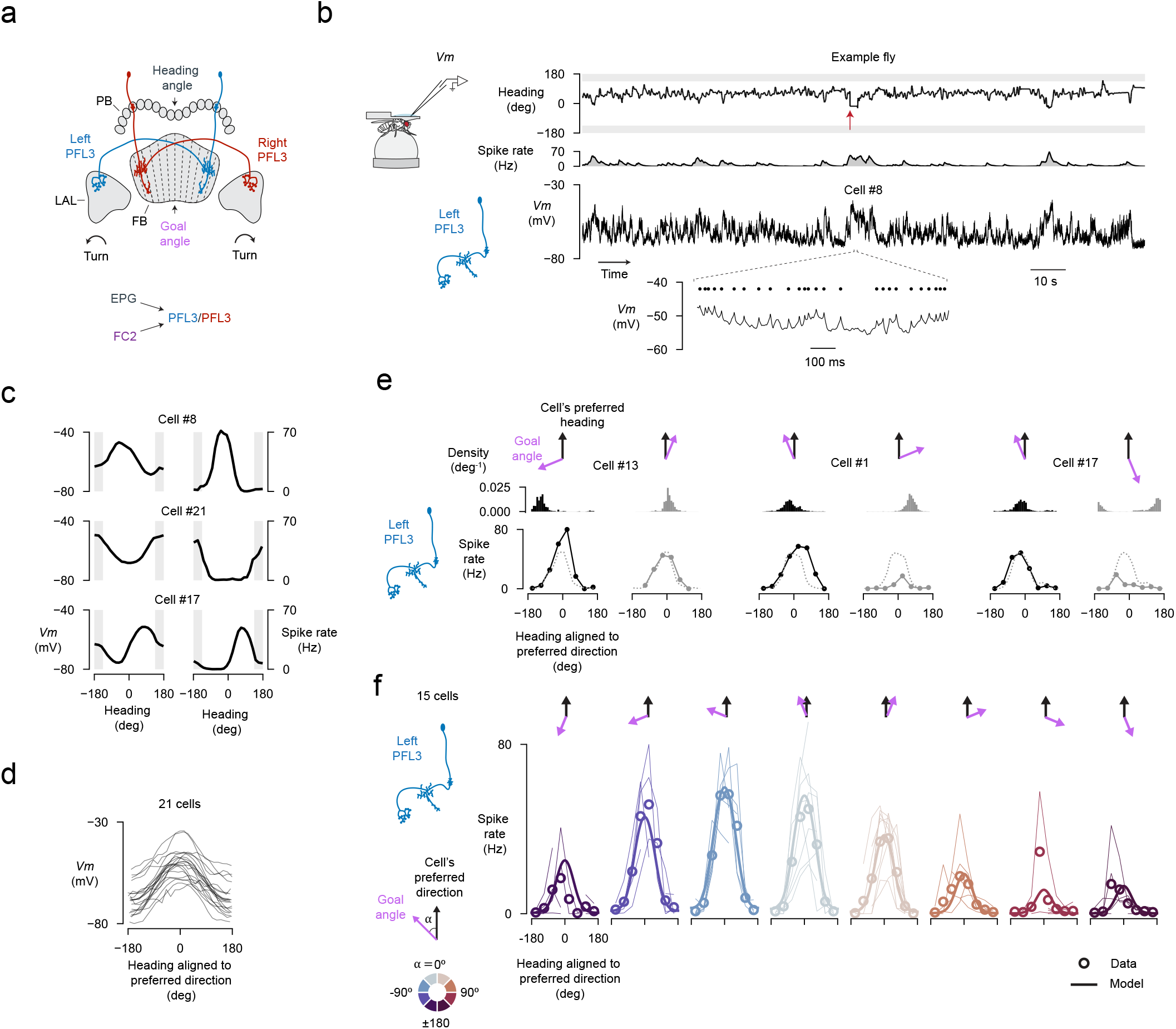
The spike-rate of PFL3 neurons shows conjunctive tuning to heading and goal angles. **a**, Two schematic PFL3 neurons. Left and right PFL3 neurons are labeled based on whether they innervate the left or right lateral accessory lobe. PFL3 neurons receive inputs from EPG neurons in the protocerebral bridge1^4,15^ (which can bring in a heading angle signal) and FC2 neurons in the fan-shaped body^14,15^ (which can bring in a goal angle signal). **b**, PFL3 patch-clamp data from a fly performing menotaxis. Top row: fly’s heading relative to the bar over time. Red arrow marks a 90° bar jump (in open-loop). Third row: membrane potential (*Vm*) recorded at the soma. Bottom row: expanded view of the membrane potential. Spikes are marked with black dots. **c**, Left: *Vm* (spikes removed) tuning curves to heading for three example cells. Right: Spike rate tuning curves from the same three cells. **d**, *Vm* (spikes removed) tuning curves for all PFL3 neurons in our population, aligned to each cell’s preferred-heading direction. **e**, Tuning curves for three example left-PFL3 neurons binned according to the angular difference between the fly’s goal angle and the cell’s preferred-heading direction. The tuning curve amplitudes are greater when the fly’s goal is to the left of the cell’s preferred direction (black) compared to when it is to the right (grey). Dashed line shows the tuning curve using data from the entire recording. Top row: histogram of behavioral heading angles expressed by the fly (aligned to the cell’s preferred direction) in association with the spike-rate tuning curves shown below. Intermittent bar jumps helped us sample heading angles outside of the relatively narrow heading distribution experienced by a fly performing menotaxis. **f**, Population-averaged, spike-rate tuning curves to heading, parsed by the flies’ goal angle. Each column represents a different bin of goal angles relative to the cell’s preferred direction. Thin lines: individual-cell tuning curves. Open circles: mean across cells. Thick lines: model fit (Methods).

To test whether PFL3 neurons might combine heading- and goal-related information, we conducted whole-cell patch clamp recordings from these cells while flies were performing menotaxis (Fig. 3a-b, Extended Data Figs. 8, 9a-c). We interspersed ±90° virtual rotations (Fig. 3b, red arrow), using the same virtual reality environment and protocol as in our imaging experiments. We identified many menotaxis bouts in these data, which allowed us to assign a behavioural goal angle—defined as the fly’s mean heading angle during a menotaxis bout—to all analyzed moments in a trajectory (Extended Data Fig. 2a-e, Methods).

Analyzing full recording sessions (which could be up to two-hours long), we generated membrane potential (*Vm*) and spike-rate tuning curves to the fly’s heading. Both the *Vm* and spike rate of PFL3 neurons were strongly tuned to heading, with different cells showing different preferred-heading directions (Fig. 3c, Extended Data Fig. 9d-e). In particular, the *Vm* displayed an approximately sinusoidal tuning to the fly’s heading (Fig. 3d, Extended Data Fig. 9d). These results are consistent with PFL3 neurons receiving heading input from the EPG and Δ7 neurons in the bridge.

To test whether the activity of PFL3 neurons depends on the fly’s goal angle as well, we re-plotted the heading-tuning curves of PFL3 neurons parsed by the fly’s goal angle. For similar heading directions, the spiking activity of PFL3 neurons varied markedly depending on the fly’s goal (Fig. 3e, Extended Data Figs. 10,11a). Specifically, the spike-rate tuning curves from left PFL3 neurons had strongly reduced amplitudes when the fly’s goal was to the right of the cell’s preferred-heading direction (Fig. 3e, Extended Data Fig. 11a). Because individual flies typically adopted only a few goal angles during an experiment, we averaged the tuning curves across all flies to provide a cell-averaged estimate for how the goal angle modulates heading-tuning in PFL3 neurons (Fig. 3f). On average, left PFL3 neurons expressed tuning curves of largest amplitude when the fly’s goal was approximately 50° to 70° to the left of the cell’s preferred-heading direction (Fig. 3f) and we observed the opposite trend in right PFL3 neurons (Extended Data Fig. 11b, bottom).

### Controlling for walking statistics

Because flies that perform menotaxis show different walking statistics depending on their angular orientation relative to the goal^13^––flies walk forward faster when aligned with their goal, for example––a neuron that is modulated by the fly’s locomotor statistics alone could yield results akin to those observed in PFL3 cells. Importantly, however, when we analyzed moments when the animals stood still, or nearly still, we observed a qualitatively similar scaling in the amplitude of PFL3 tuning curves (Extended Data Fig. 12), indicating that PFL3 goal direction modulation is not a simple consequence of the fly’s walking dynamics, but is more likely to be generated by FC2 inputs or a similar input signal.

### A model for single-cell PFL3 responses

The conjunctive tuning of PFL3 neurons to heading and goal angles (Fig. 3f), along with the shape of the spike-rate vs. *Vm* response function (Extended Data Fig. 13c), led us to formulate a descriptive model of the single-cell tuning properties of PFL3 neurons (Extended Data Fig. 13; Methods). Specifically, we modeled the PFL3 spike rate as a nonlinear function of the sum of two sinusoids. One sinusoid represents the EPG/Δ7 input in the bridge, which is expected and observed (Fig. 3c, Extended Data Fig. 9d) to show sinusoidal tuning to heading^4^. The second sinusoid represents the goal input in the fan-shaped body, which also appears to be sinusoidal (Extended Data Fig. 13d). We thus model the activity of a single PFL3 neuron as *f* (cos(*H* − *φ*) + *d*cos(*G* − *θ*)), where *H* is the fly’s heading angle, *G* is the its goal angle, and *φ* and *θ* are the preferred heading and goal angles, respectively, for the PFL3 cell being modeled. The parameter *d* accounts for the relative strengths of the heading- and goal-dependent inputs. The form of the nonlinear function *f* was obtained from the firing rate versus *Vm* curves of actual PFL3 neurons (Extended Data Fig. 13b-c, Methods). We fit this model to the data in Fig. 3f. Because the curves in this figure have been shifted by the preferred heading angle *φ*, the fit only depends on the difference *θ* − *φ*, which we take, for now, to be the same for all cells, along with *d* and the three parameters describing the function *f* (Methods). This model captures the heading- and goal-dependences of spike-rate tuning curves from PFL3 cells quite well (Fig. 3f, R^2^=0.91).

### A circuit model for goal-directed steering

To gain intuition for how PFL3 neurons with the above single-cell properties could direct turning toward a goal, consider a scenario consisting of two PFL3 neurons (one left and one right) that project to a common fan-shaped body column. Because these two cells receive shared inputs in the fan-shaped body (Extended Data Fig. 7e-h), any differences in their activity would be determined entirely by their heading input from the bridge, which is expected to be different because their preferred-heading directions are offset from one another (Fig. 4a, red and blue arrows). If the fly’s heading is aligned with the right cell’s preferred-heading angle, the activity of the right cell will be greater than that of the left cell. This would create an asymmetry in the left and right LAL activity appropriate for directing a rightward turn (Fig. 4a, bottom). The opposite would be true if the fly was aligned with left cell’s preferred heading. In this simple scenario, a fly would orient along a fixed angle, midway between the preferred heading angles of the left-right pair (purple arrow). However, with only two PFL3 neurons at its disposal, a fly would be limited to a single, inflexible, goal angle. This limitation is removed by considering a model of the full PFL3 population. Building on previous computational studies^18,27–29^ and applying insights from our FC2 experiments, PFL3 recordings, and single-cell PFL3 model, we developed such a circuit model for how PFL3 neurons enable goal-directed steering.

**Fig. 4.**
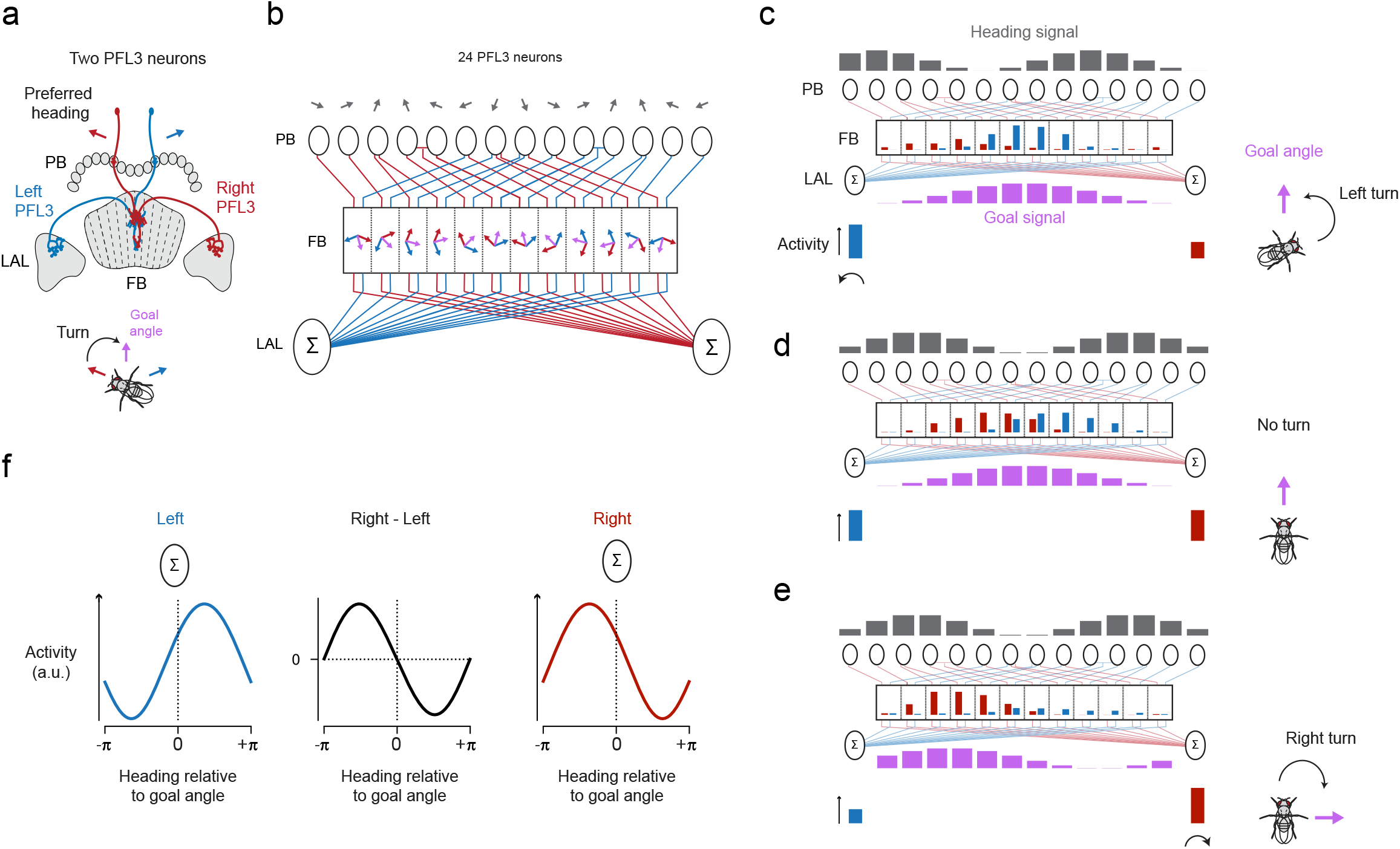
Model for how PFL3 neurons compare heading and goal angles to generate a steering signal. **a**, Schematic of two PFL3 neurons with offset preferred-heading directions (red and blue arrows) that they inherit in the bridge. The two cells project to a common column in the fan-shaped body. These two PFL3 cells could lead a fly to behaviourally stabilize a single allocentric goal angle, midway between the preferred heading angles of the two cells (purple arrow in diagram below). **b**, Wiring diagram of all 24 PFL3 neurons in the fly brain^15^. Each grey arrow represents the preferred-heading angle that a PFL3 neuron innervating a given glomerulus of the protocerebral bridge (PB) is expected to inherit from presynaptic heading-sensitive EPG/Δ7 neurons in that glomerulus (see Extended Data Fig. 6). Blue and red arrows represent the bridge-inherited, preferred-heading direction angle *φ* of the left and right PFL3 neurons that innervate a given column in the fan-shaped body (FB). Purple arrows represent each column’s preferred goal angle θ. Angles are measured positive counterclockwise with zero pointing straight downward. **c**, Example heading and goal input bumps to the PFL3 population and the predicted output signal from individual PFL3 neurons and the PFL3 population. The neural signals in the schematic would be evident in the brain in apply to the situation depicted by the fly on the right. Dark grey bar plots: the spatial activity pattern of the heading inputs to PFL3 cells in the protocerebral bridge. Purple bar plots: the spatial activity pattern of goal (FC2) inputs to PFL3 cells in the fan-shaped body. Red and blue bar plots in fan-shaped body represent the activity of individual PFL3 neurons, determined by a nonlinear function of their summed bridge and fan-shaped body inputs. Red and blue bar plots below the sigma symbol: summed activity for left and right PFL3 neurons in the lateral accessory lobe (LAL). **d-e**, Same as panel c but for different heading and goal angles. **f**, Model-predicted, population-level activity in the left and right LAL (blue and red curves) and predicted turning signal (right-minus-left LAL activity, black curve).

This model is based on the single-cell fit described in the previous section, but rather than fitting the difference in preferred heading and goal angles, *φ* and *θ*, we determine these angles separately and independently for each PFL3 on the basis of connectomics data^15^ (Fig. 4b). All other parameters (*d* and the parameter describing *f*) are taken from the fit in Figure 3f. As in the two-cell scenario described in the previous paragraph, each fan-shaped body column is innervated by two PFL3 neurons, one projecting an axon to the right LAL and the other to the left LAL. Critically, pairs of PFL3 neurons that innervate the same column in the fan-shaped body receive inputs from different glomeruli in the protocerebral bridge (Fig. 4b). Each bridge glomerulus can be assigned an angle based on the direction the fly would be heading if the EPG or Δ7 bumps expressed their maximum activity within that glomerulus^4^ (Fig. 4b, grey arrows, Extended Data Fig. 6). The preferred-heading angles of PFL3 neurons can be inferred from these bridge angles on the basis of their projections from the bridge to the fan-shaped body (Extended Data Fig. 6a-c). The red and blue arrows within each fan-shaped body column in Figure. 4b indicate the preferred heading angle assignments for the left (blue) and right (red) PFL3 neurons innervating that column (for values see Methods). The preferred-goal angles are obtained by dividing the full 360° spanned by the columns of the of the fan-shaped body into twelve equally spaced values (Fig. 4b, purple arrows). Collectively, this anatomy results in an array of twelve left/right PFL3-pairs with preferred heading and preferred goal angles that span azimuthal space.

The operation of the model for three different heading-goal relationships is shown in Figure 4c-e. The grey bars at the top of these figures show the heading-related input received by the PFL3 cells within each glomerulus of the protocerebral bridge. The height of each bar is proportional to the cosine of the angle between the heading direction of the fly (shown to the right of the figures) and the corresponding (grey) preferred-heading arrow at the top of Figure 4b. These heading signals are conveyed to the columns of the fan-shaped body through the connections depicted by faint red and blue lines and in Figure 4b. The goal-related FC2 input for each fan-shaped-body column is indicated by the purple bars at the bottom of the figures. The height of each bar is proportional to the cosine of the angle between the fly’s goal (purple arrow shown to the right of the figures) and the purple arrow within each column in Figure 4b. Red and blue bars within the columns shown in Figure 4c-e show the resulting PFL3 activities, which are given by the sum of the inputs depicted by the appropriate grey and purple bars passed through the nonlinear PFL3 input-output response function (*f*).

The full model operates in a manner that is a generalization of our description of Figure 4a. When the heading and goal angles align (Fig. 4d), the activity of left and right PFL3 cells does not match within every column, but it does match overall. As a result, the left and right LAL signals, which are given by sums over all of the left or right PFL3 neurons, are equal (Fig. 4d). We assume that the turning signal generated by the PFL3 cells is the difference between the right and left LAL activities. Thus, when the heading and the goal align, there is no net turning signal. If the fly is headed to the right of the same goal (Fig. 4c), the goal input does not change from the previous example, but the heading signal does. This breaks the left-right balance, making the total activity of the left PFL3 cells greater than that of the right PFL3 cells. The resulting imbalance in the left and right LAL signals then generates a turn signal to the left. If, on the other hand, the goal direction changes (Fig. 4e), the change in the goal signal (purple bars) breaks that balance, resulting, in this case, in greater total right than left PFL3 activity in the LAL, producing a rightward turning signal.

Our model predicts the summed PFL3 activity in the left and right LALs as a function of the fly’s heading relative to is goal angle (Fig. 4f, red and blue curves). The difference between these two signals corresponds to the steering signal that we expect flies to use in stabilizing their trajectory during menotaxis (Fig. 4f, black curve). This predicted turning signal has a sinusoidal shape in close agreement with previous behavioral measurements in menotaxis^13^.

If the difference between the right and left summed LAL activities controls turning, the fly will maintain a heading defined by the angle where the turning signal is zero and its slope is negative (the zero crossing at the center of the middle panel in Fig. 4f). In the model, we find that this ‘zero’ heading direction is equal to the phase of the goal signal, on average, but with a standard deviation of 2° and a maximum deviation of 3.3° across the full range of goal directions (Extended Data Fig. 13f,g). These small ‘errors’ arise solely from the fact that two of the PFL3 cells innervate two glomeruli of the protocerebral bridge (Fig. 4b, Extended Data Fig. 6)––if they innervated a single glomerulus like all the other PFL3 cells, the steering would match the goal direction perfectly––making this a curious feature of the anatomy.

### PFL3 physiology supports the model

To test the predictions of the model, we performed two-photon calcium imaging of the axon terminals of PFL3 neurons in the right and left LALs (Fig. 5a). Transient increases in the right-minus-left GCaMP signal were, on average, followed by an increase in rightward turning (with ∼100 ms latency) and vice versa (Fig. 5b-c), as expected if a LAL asymmetry in PFL3 activity acts to promote turning in the appropriate direction. We plotted PFL3 GCaMP activity in the left and right LAL, separately, as well as the difference between these two signals, as a function of the fly’s heading relative to the goal. The left and right PFL3 curves (Fig. 5d, top two rows)–– which peaked at headings approximately ±70° from the fly’s goal––alongside the difference between these two signals (Fig. 5d, bottom row), match our expectations from the model (Fig. 4f); the shapes of the model curves are quite close to those of the data curves, but there are small shifts between the model and data curves along the horizontal axis (Extended Data Fig. 13e). The experimentally measured difference curve appeared to be sinusoidally shaped, as expected both from our model (Fig. 4f) and from our previous behavioral measurements^13^. Finally, to test whether experimentally activating PFL3 cells in the LAL could cause flies to turn, we optogenetically stimulated either the left or right LAL of flies that co-expressed CsChrimson and jGCaMP7f in PFL3 neurons (Fig. 5e). Co-expressing GCaMP in the same cells allowed us to calibrate our stimulation levels to elicit a desired level of GCaMP signal. We observed an increase in ipsilateral turning during the 2 s stimulation period, which was not observed in control flies that did not express CsChrimson (Fig. 5f-h). In addition, when we performed the same experiment with PFL1 neurons^14,35,44^–– a morphologically similar cell type with different connectivity^14,15^––we did not observe an increase turning velocity during stimulation (Fig. 5g-h), even though the LAL GCaMP signal indicated that PFL1 neurons were strongly activated during these experiments. The PFL1 result shows that ipsilateral turning is not an inevitable outcome of strong asymmetric stimulation of any cell class in the LAL.

**Fig. 5.**
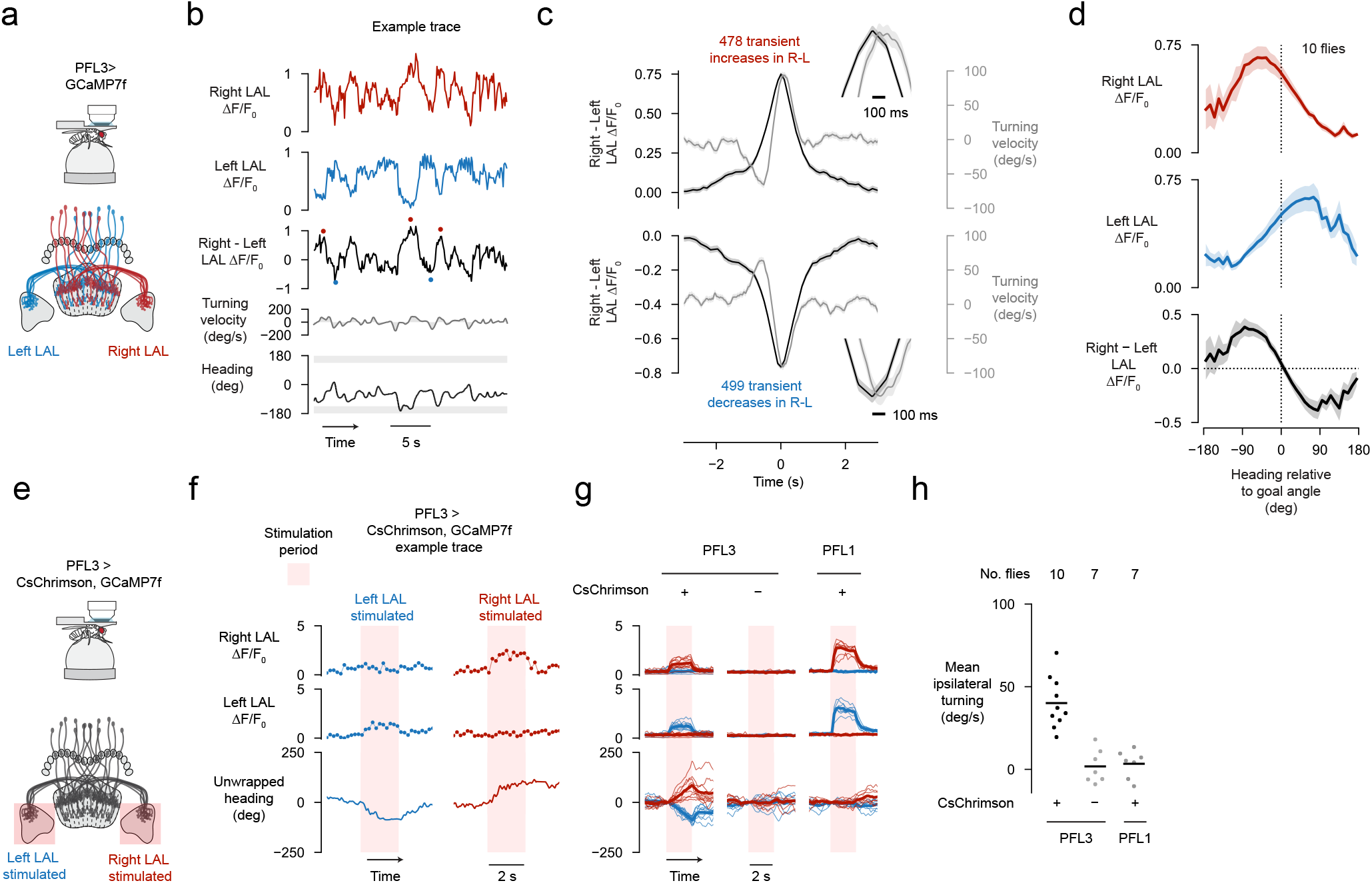
Imaging and perturbing PFL3 activity in the lateral accessory lobes supports the model. **a**, Two-photon calcium imaging of the lateral accessory lobe (LAL) of flies expressing jGCaMP7f in PFL3 neurons labeled by split-Gal4 line 57C10-AD ∩ VT037220-DBD. **b**, Example timeseries of GCaMP imaging data. In the third row, red dots mark transient increases in the right - left (R-L) ΔF/F0 signal and blue dots mark transient decreases. **c**, The flies’ turning velocity (grey) and R-L signal (black) aligned to transient increases (top) or decreases (bottom) in the R-L signal. Insets show that the peak in the R-L asymmetry precedes the peak in turning velocity by ∼100 ms (see Methods). Mean ± s.e.m. across transients is shown (from 10 flies). **d**, LAL activity plotted as a function of the fly’s heading relative to its goal angle. Mean ± s.e.m. across flies is shown. **e**, Stimulation of PFL3 cells in either left or right LAL while simultaneously performing calcium imaging from the same cells. We used flies that co-expressed CsChrimson and jGCaMP7f in PFL3 neurons labeled by split-Gal4 line VT000355-AD ∩ VT037220-DBD. **f**, Left: example trial where we stimulated the left LAL. Bottom row, unwrapped heading zeroed at onset of stimulation. A decrease in the unwrapped heading signal means the fly turned left. Right: example trial with the right LAL stimulated. **g**, Fly-averaged GCaMP and turn signals (thin lines) for left (blue) and right (red) LAL stimulation of PFL3 or PFL1 cells. Thick line shows the average across flies. **h**, Mean ipsilateral (relative to the stimulation side) turning velocity during the 2 s stimulation period. Dots show the mean for individual flies and lines indicate the mean across flies. PFL3 CsChrimson flies have a greater ipsilateral turning velocity than non CsChrimson PFL3 flies (p=1.93×10-5, Welch’s two-sided t-test). PFL1 Chrimson flies show no change ipsilateral turning velocity relative to controls (p=0.76, Welch’s two-sided t-test).

## Discussion

When navigating toward a visible object, or performing a behaviour like phonotaxis, sensori-motor transformations within an egocentric reference frame are sufficient; if the object is sensed to be on the left, one should turn to the left. However, if an animal wishes to orient toward a remembered direction or location, the underlying computations become greatly simplified if the animal can employ a common allocentric reference frame for signaling variables of interest. Prior work had identified bumps of calcium activity that track a fly’s allocentric heading and traveling angles^2,4,5,13^. For a fly to navigate in a desired direction its brain needs to track not only spatial variables relating the fly’s *current* state, but also variables that signal the fly’s *desired* state in the same allocentric reference frame. We describe here an activity bump, carried by FC2 neurons, that can signal the fly’s allocentric goal direction (Figs. 1-2). We also described how PFL3 neurons combine allocentric heading and goal signals to generate an egocentric turning signal (Figs. 3-5).

Our FC2 experiments are consistent with these cells *communicating* an angular goal to downstream circuits. On the other hand, the FC2 calcium signal need not be *storing* the fly’s goal. Indeed, during menotaxis, we noticed instances when the FC2 calcium signal drifted in the fan-shaped body with changes in the fly’s heading, but the fly’s goal, on a longer timescale, seemed to remain unchanged (Extended Data Fig. 3, teal arrow). One interpretation of this result is that long-term angular goals are either stored in a latent, molecular signal in FC2 cells that is not always evident in calcium, or alternatively, in other cells or synapses with a variable ability to drive FC2 cells. It has been argued, via free-behaviour experiments, that sparsely stimulating a set of columnar neurons upstream of FC2 cells can induce flies to walk along certain directions^45^. In addition, the fan-shaped body may encode multiple ‘potential’ goals, with the actual goal chosen from this set in a state-dependent manner. Thus, the FC2 calcium signal might be best viewed as a conduit between long-term navigational goals and the central-complex’s pre-motor output. Given that FC2 neurons are only two synapses away from descending neurons, it is perhaps not surprising that their activity seems to more closely reflect the fly’s desired behaviour on a shorter timescale.

For the central complex to control behaviour, allocentric signals need to be converted to egocentric signals appropriate for the motor system. Our work provides a physiological account for how PFL3 neurons accomplish this coordinate transformation. Specifically, individual PFL3 neurons combine heading and goal direction inputs to produce, as a population, an error signal that informs the motor system whether to turn left or right. Mathematically, this circuit can be considered to be projecting a vector that encodes the fly’s allocentric goal angle––signaled by the position of the FC2 bump in the fan-shaped body––onto two axes linked to the fly’s heading direction. One axis represents the fly’s heading angle rotated clockwise and the second axis represents the fly’s heading angle rotated counter-clockwise by the same amount. The difference between the projections of the goal vector onto these axes indicates how much, and in which direction, the fly should turn to orient itself toward the goal angle (Extended Data Fig. 14).

Studies in mammals have identified neurons that track an animal’s egocentric bearing to a point in space, an object in the local environment, or a goal location^10–12,46^. For instance, the bat hippocampus houses neurons that fire maximally when a landing perch is at a specific angle relative to the bat’s current heading^10^. Analogous to the bat neurons, the summed population activity of PFL3 neurons in the left or right LAL is tuned to specific heading angle relative to the fly’s goal (Fig. 5d red and blue curves). This observation suggests that the computations implemented by the PFL3 system may ultimately find analogies in the mammalian brain.

Neurons that are anatomically downstream of PFL3 cells in the LAL, including descending neurons that project to the ventral nerve cord^27^, are poised to sum the output of the left PFL3 cells and, separately, the right PFL3 cells. Although our PFL3 imaging and stimulation experiments indicate that asymmetries between these two signals can impact steering behaviour (Fig. 5), exactly how downstream motor centers interpret PFL3 signals remains to be determined. The effect of PFL3 output on locomotor behaviour could be gated by the fly’s internal state or the context in which the fly finds itself. Moreover, many other signals that could affect turning converge onto the known, locomotion-related descending neurons^14,27^. Thus, the turning signal we have discovered here is more likely to affect the probability of turning in a certain direction rather than driving a deterministic, short-latency motor response. It has been similarly suggested that such a PFL3 signal should impact the fly’s locomotory “policy”^29^.

Modeling work on a visually-guided learning task^29^ argued that an angular goal signal in the fan-shaped body might take an arbitrary shape, rather than being constrained to a profile with a single peak. When stimulating FC2 neurons, we noticed a decrease in activity in non-stimulated FC2 columns (Extended Data Fig. 5c-d), suggesting the presence of feedback inhibition ensuring that only a single bump of activity or goal is relayed to PFL3 neurons at any one time. The modeling ideas could be reconciled with our FC2 data if the storage of goal information is physiologically separated from its comparison with the fly’s heading angle. That is, the goal-memory signal could, for example, exist in a set of synaptic weights to the FC2 system^29^, with a winner-take-all circuit converting this potentially more complex input into a single peak of calcium activity that is ultimately relayed to the PFL3 systems.

Although our experiments were limited to menotaxis––a behaviour that requires the animal to determine in which direction to walk, and not necessarily how far––we speculate that the FC2-PFL3 circuit also functions to regulate turning when the animal is navigating toward a location in 2D space. A previous study modeled how the fan-shaped body could produce signals to steer a bee back to its nest^18^. In this model, PFL3-like neurons^3^ receive direct inputs from a columnar array of home-vector neurons, with the phase and amplitude of the array’s sinusoidal signal representing the allocentric angle and distance to the nest, respectively. In this scheme, as the bee gets closer to its nest and the home vector gets shorter, the amplitude of the neuronal homing/goal signals gets smaller. This means that a turning signal derived from such neurons will become noisier near the goal. One possibility, hinted at by our data, is that 2D goal vectors are, at some point, split into separate distance and direction signals, with the goal angle, encoded by a distance-independent bump of activity with a high signal-to-noise ratio, guiding turning. Using a purely angular comparison as the final step in deciding whether to turn right or left could overcome issues with noise, and might be a primitive used in many navigational behaviours.

## Acknowledgements

We thank members of the Maimon laboratory for helpful discussions. We thank Jonathan Green, Cheng Lyu and Itzel Ishida for feedback on the manuscript. We thank Sachin Sethi and Cheng Lyu for technical advice on patch-clamp experiments. We thank Jazz Weisman for the design and fabrication of various 3D printed parts. We thank Tobias Nöbauer for help setting up two-photon light paths. We thank Jim Petrillo for the fabrication of fly-tethering plates. We thank Georg Jaindl for software advice for simultaneous imaging and stimulation experiments. Stocks obtained from the Bloomington Drosophila Stock Center (NIH P40OD018537) were used for this study. Research reported in this publication was supported by a Brain Initiative grant from the National Institute of Neurological Disorders and Stroke (R01NS104934) to G.M. L.A. was supported by the Simons, Gatsby and Kavli Foundations and by NSF NeuroNex Award DBI-1707398. G.M. is a Howard Hughes Medical Institute Investigator.

## Author contributions

P.M.P. and G.M. conceived of the project. P.M.P. performed the experiments and analyzed the data. P.M.P., L.A., and G.M. jointly interpreted the data. L.A. developed and implemented the formal models. P.M.P., L.A., and G.M. wrote the paper together.

## Author information

The authors declare no competing financial interests.

---

**Extended Data Fig. 1.**
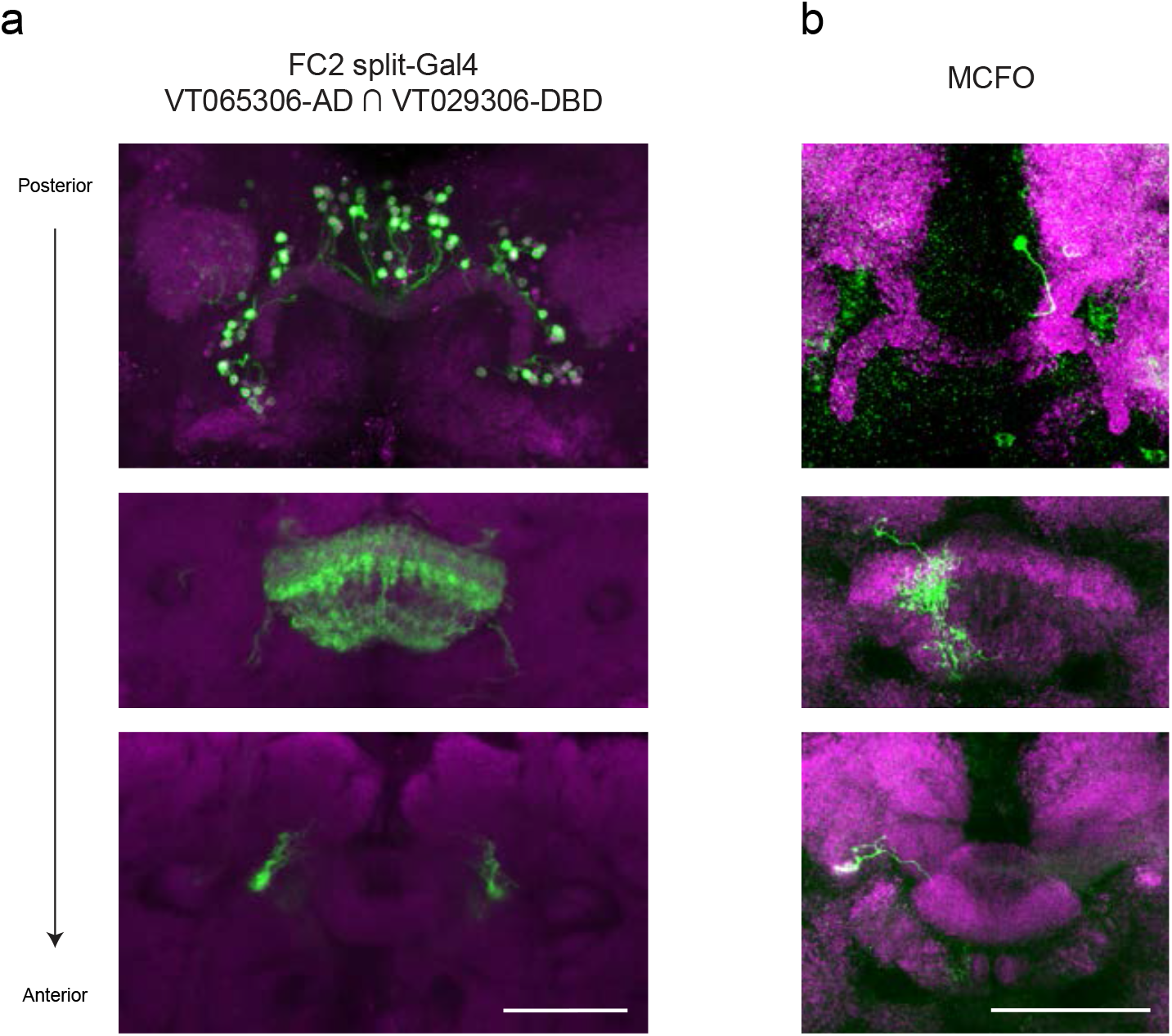
FC2 split-Gal4 line characterization. **a**, GFP expression driven by split-Gal4 line VT065306-AD∩VT029306-DBD (green) and anti-nc82 neuropil stain (magenta). Each panel shows an average z-projection. Top: the number of GFP positive somas, roughly 70 to 100, is comparable to the 88 FC2 neurons identified in the hemibrain^14^. While the GFP expression in this line suggests that it is selective for crepine projecting neurons with FC2-like anatomy, it is possible that there are some non-FC2 central complex neurons labeled by the line as well. Middle: fan-shaped body. Bottom: Crepine. Each FC2 neuron projects unilaterally to the crepine, a symmetric structure that flanks the central complex and is situated dorsal to the lateral accessory lobes. **b**, Multicolor flip-out of an FC2 neuron labeled VT065306-AD∩VT029306-DBD. The innervation pattern in the fan-shaped body is consistent with the FC2B or FC2C subtypes.

**Extended Data Fig. 2.**
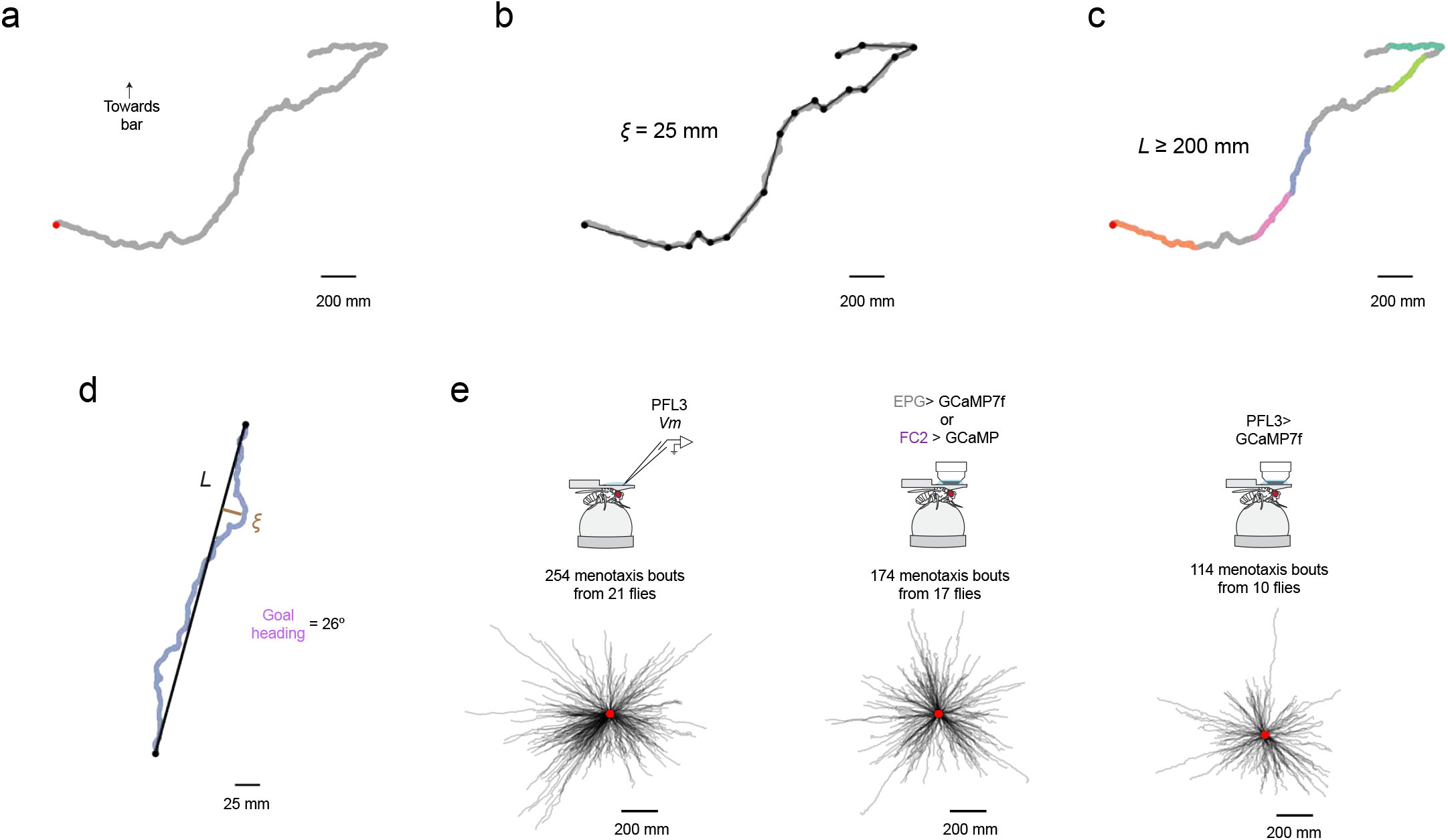
Using the fly’s virtual 2D trajectory to analyze menotaxis behaviour. **a**, Example virtual 2D trajectory of a fly performing menotaxis (during a PFL3 patch-clamp recording). Red dot marks the start of the trajectory. **b**, Ramer-Douglas-Peucker algorithm reduces the number of *x*,*y* coordinates in the trajectory using the parameter S, the maximum allowed distance between the simplified and original trajectories. Black dots show the simplified coordinates. **c**, The fly’s displacement between each *x*,*y* point of the simplified trajectory, *L*, is computed. Segments of the fly’s trajectory where *L* ≥ 200 mm were considered “menotaxis bouts” and thus further analyzed (colored portions of the trajectory). **d**, An example menotaxis bout from trajectory in panel **c**. The fly’s goal angle is defined as the fly’s mean heading direction during the bout, excluding timepoints when the fly is standing still. **e**, All menotaxis bouts from flies used in this paper. Left: PFL3 patch-clamp dataset (related to Fig. 3). Middle: EPG and FC2 imaging dataset (related to Fig. 1). Right: PFL3 LAL imaging dataset (related to Fig. 5d).

**Extended Data Fig. 3.**
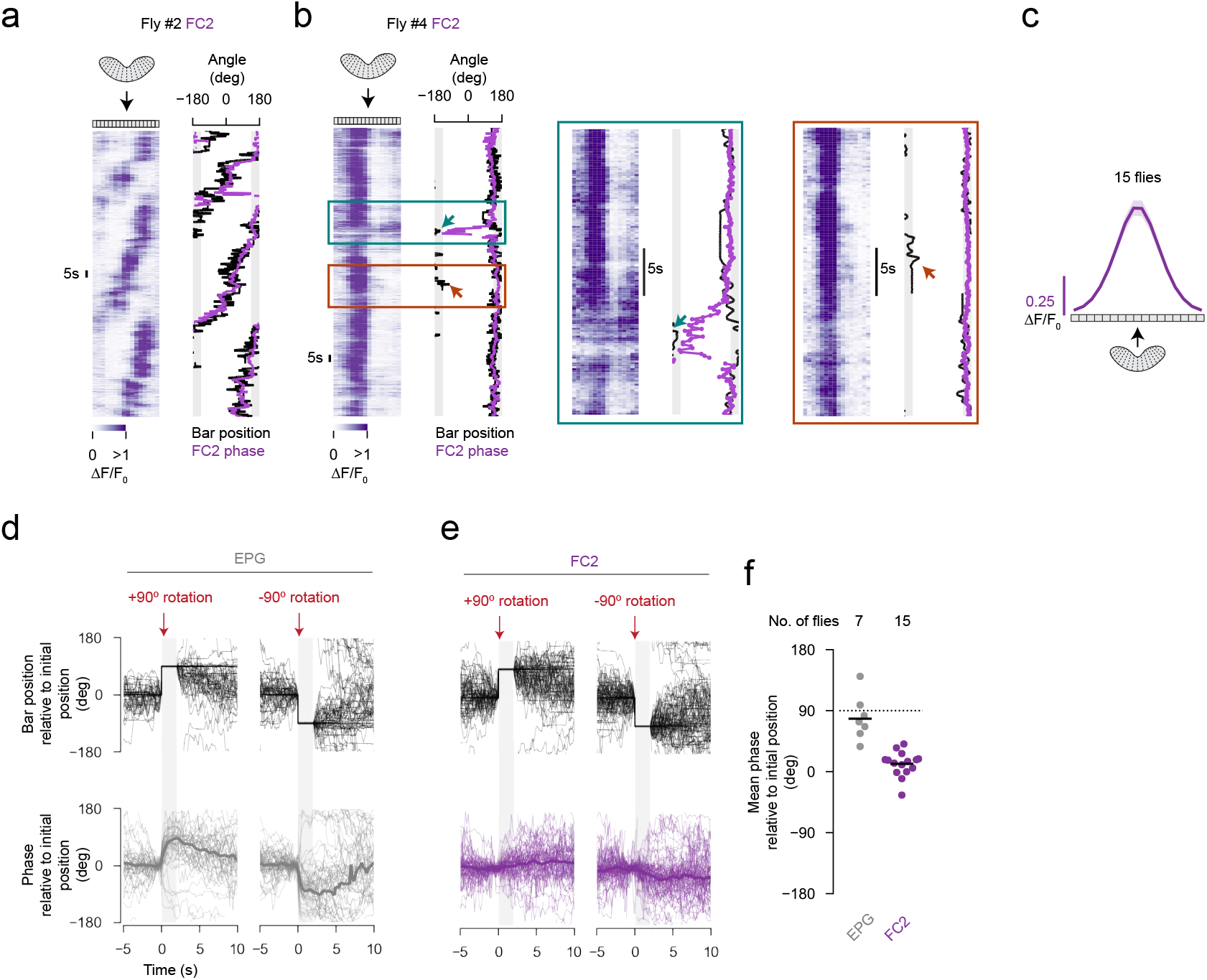
Example traces of the FC2 bump alongside EPG and FC2 phase responses to virtual rotations without selecting trials based on the fly’s behaviour. **a**, Example FC2 ΔF/F0 signal and behavioral traces from a fly that was rotating in time and not stabilizing a consistent heading direction. This trace highlights that the FC2 phase can be well-estimated during moments where our algorithm would not detect that the fly is performing menotaxis. Our working model for such moments is that central-complex independent circuitry is driving turning and that the FC2 phase passively follows the fly’s heading, until it is needed to govern behavioural output. **b**, Left: Example FC2 ΔF/F0 signal and behavioral traces from a fly that occasionally deviated from its goal angle. The teal arrow marks a moment when the FC2 phase did not remain stable, but the fly nonetheless returned to its putative goal direction. The FC2 cells, we believe, are interposed between phasors that store medium-to-long-term vector memories and the ability of the central complex to use such vector memories to impact locomotor turns. What might have happened in the moment marked in teal is that inputs other than the longer-term menotaxis goal input to the FC2 system briefly dominated—or no inputs were dominant for a brief moment—which led the FC2 phase to drift. However, once the fly re-entered the menotaxis behavioral state and wished progress forward, the FC2 phase locked back in to that angle, communicating it to the PFL3 population to guide steering. The red arrow marks an occasion when the FC2 phase remained stable throughout a brief deviation in heading direction. Right: expanded view of time period marked by teal box and red box. **c**, Phase-nulled FC2 activity averaged across all flies. Mean ± s.e.m. across flies is shown. **d**, Individual ±90° rotation trials for 93 trials from 7 flies in which we imaged EPG neurons. In this case, we did not require for a trial to occur within a menotaxis bout (see Methods) or require that the fly return within 45° from its heading before the bar jump. Thick lines show mean across flies. **e**, Same as panel d but for 141 trials from 15 flies in which we imaged FC2 neurons. Note that, on average, the FC2 phase slowly drifts away from its initial position. This small drift is likely due to trials where the fly’s goal angle genuinely drifted to the fly’s new heading angle after the bar jump, which seems plausible given that on many trials analyzed here the fly did not turn so as to reorient themselves along their previous heading. **f**, Mean phase value during final 1 s of the open-loop period in panels d and e. Each dot is the mean for one fly. Horizontal lines show

**Extended Data Fig. 4.**
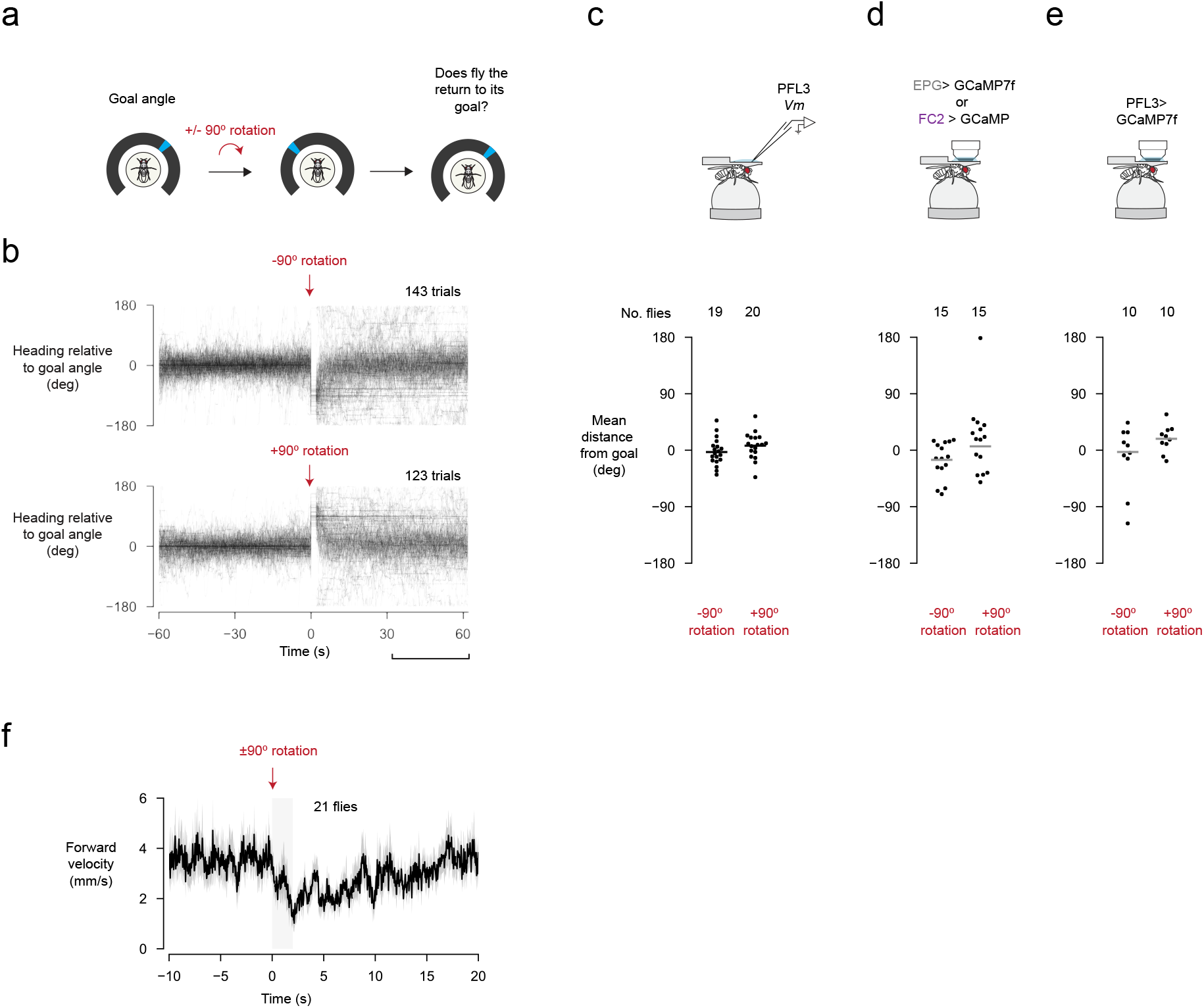
Following a virtual rotation, flies slow down and turn so as to return to their previous heading. **a**, To assess whether a fly was actively maintaining its heading direction, we virtually rotated it by discontinuously jumping the bar ±90º from its position immediately before the jump. The bar remained static at its new position for 2 s, afterwards, the fly regained closed-loop control over the bar. **b**, The fly’s heading relative to its goal angle for ±90º rotation trials from our PFL3 patch-clamp dataset. Only trials where the circular standard deviation of the fly’s heading direction during the 60 s prior to the bar jump was less than 45º were analyzed here (59% to 68% of all trials depending on the dataset). For this analysis, we defined the fly’s goal angle as its mean heading in the 30 s before the bar jump, excluding timepoints in which the fly was standing still. **c**, Mean heading relative to the fly’s goal angle during the 30 to 60 s after the bar jump. Each dot is the mean for an individual fly. Horizontal line: mean across flies. **d**, Same as panel **c** but for our EPG and FC2 imaging dataset. **e**, Same as panel c but for our PFL3 LAL imaging dataset. f, Mean forward walking velocity around the time of bar jumps. Shaded area marks the 2 s when the bar remained static. Mean ± s.e.m. across flies is shown.

**Extended Data Fig. 5.**
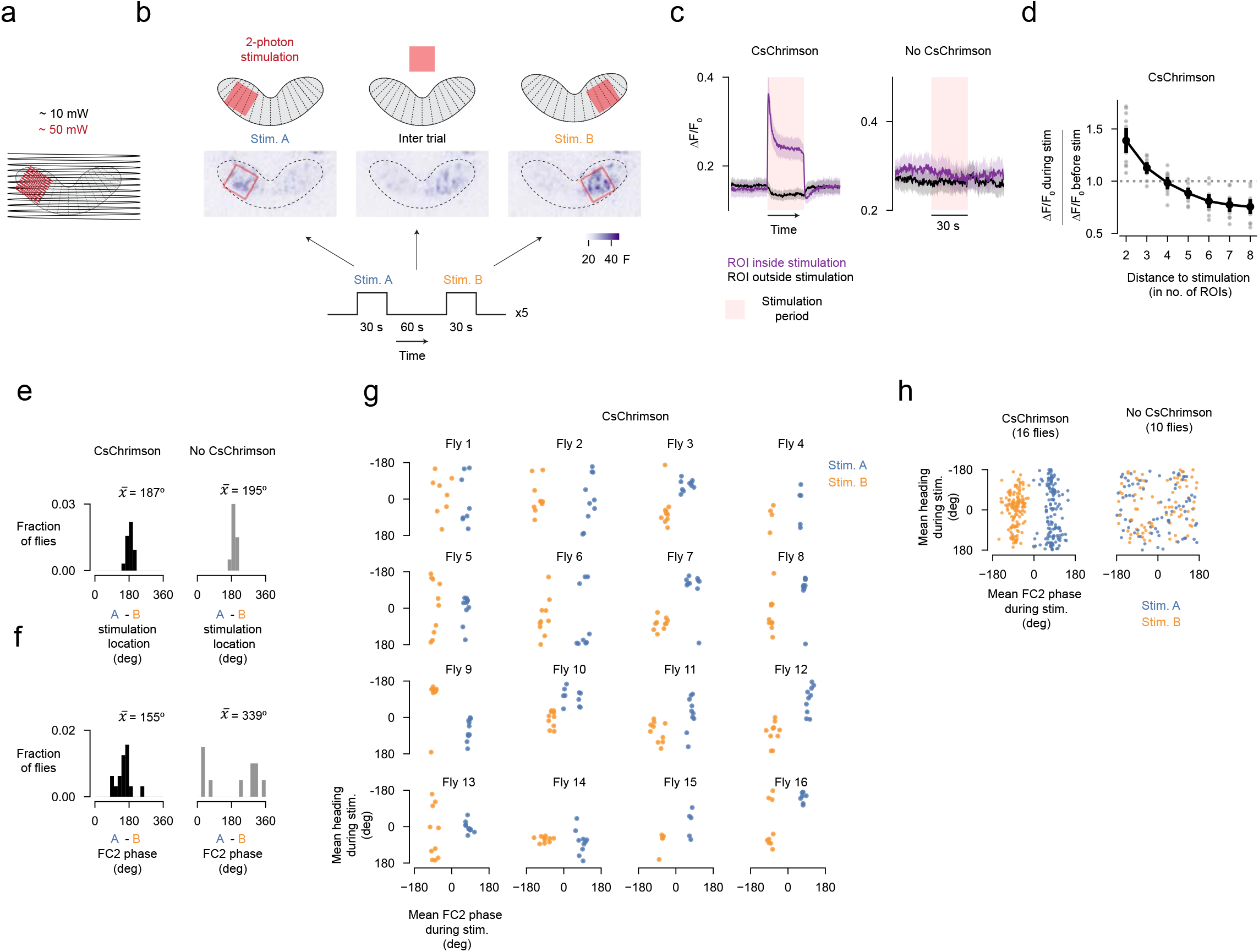
FC2 neurons in one column of the fan-shaped body inhibit FC2 neurons in distant columns and an approximately one-to-one mapping exists between the FC2 phase and the goal angle within, but not across, flies. **a**, Schematic of scan paths for the entire imaging region (black) alongside the stimulation (red) regions of interest (ROI). **b**, Trial structure for columnar stimulation. Top: 16 fan-shaped body column ROIs (regions delineated by the dotted lines) and the two stimulation ROIs (red squares). Note that the stimulation ROI can overlap with several column ROIs. Middle: average z-projection of the raw fluorescence signal during stimulation in position A (stim. A; blue), the inter-trial period and stimulation at position B (stim. B; orange). This panel shows the same example trial as in Figure 5b. **c**, Left: Mean column ROI ΔF/F0 aligned to the onset of stimulation (pink background) from flies expressing CsChrimson in FC2 neurons for ROIs that overlap with the stimulation ROI (purple) or ROIs that do not overlap with the stimulation ROI (black). Right: same as left, but for control flies that do not express CsChrimson. Mean ± s.e.m. across flies is shown. **d**, Change in non-stimulated ROI ΔF/F0 as a function of the ROI’s wrapped distance from the stimulation site for CsChrimson expressing flies. Each grey dot is the mean for an individual fly. Black dots and thick line shows mean ± s.e.m. across flies. The increase in activity of column ROIs with a distance of 2 or 3 could reflect lateral excitation or alternatively, could simply be due to neurites of stimulated neurons within the stimulation ROI extending into non-stimulated ROIs. **e**, Distribution of the estimated angular difference—assuming the fan-shaped body leftIright extent maps to 360° of azimuthal space—between stimulation location A and B for all flies (see Methods for how stimulation location angle is computed). 20° bins. **f**, Distribution of the angular difference between the mean FC2 phase position during stimulation A and B for all flies. 20° bins. **g**, Heading as a function of the FC2 phase position in the fan-shaped body for flies expressing CsChrimson in FC2 neurons. Each dot is a trial, color coded by the simulation location. A phase value of zero signifies that the FC2 bump is centered on the middle of the fan-shaped body. Note that the same phase position can be reliably associated with a similar heading direction within a fly, but not necessarily across flies (e.g. compare fly 7 to fly 9). The fact that individual flies show a unique offset between the stimulated fan-shaped body location and the stabilized behavioural heading angle is expected if the FC2/PFL3 system signals angles in the same allocentric reference frame set by the EPG heading bump. This is because the EPG bump in the ellipsoid body shows a variable fly-to-fly offset between the fly’s heading in the world and the bump-position in the brain2. **h**, Left: same data as in panel e, but all trials for all flies are shown in the same plot. Note that there is no clear relationship between phase position and bar position across flies. Right: same as left but for control flies that do not express CsChrimson in FC2 neurons.

**Extended Data Fig. 6:**
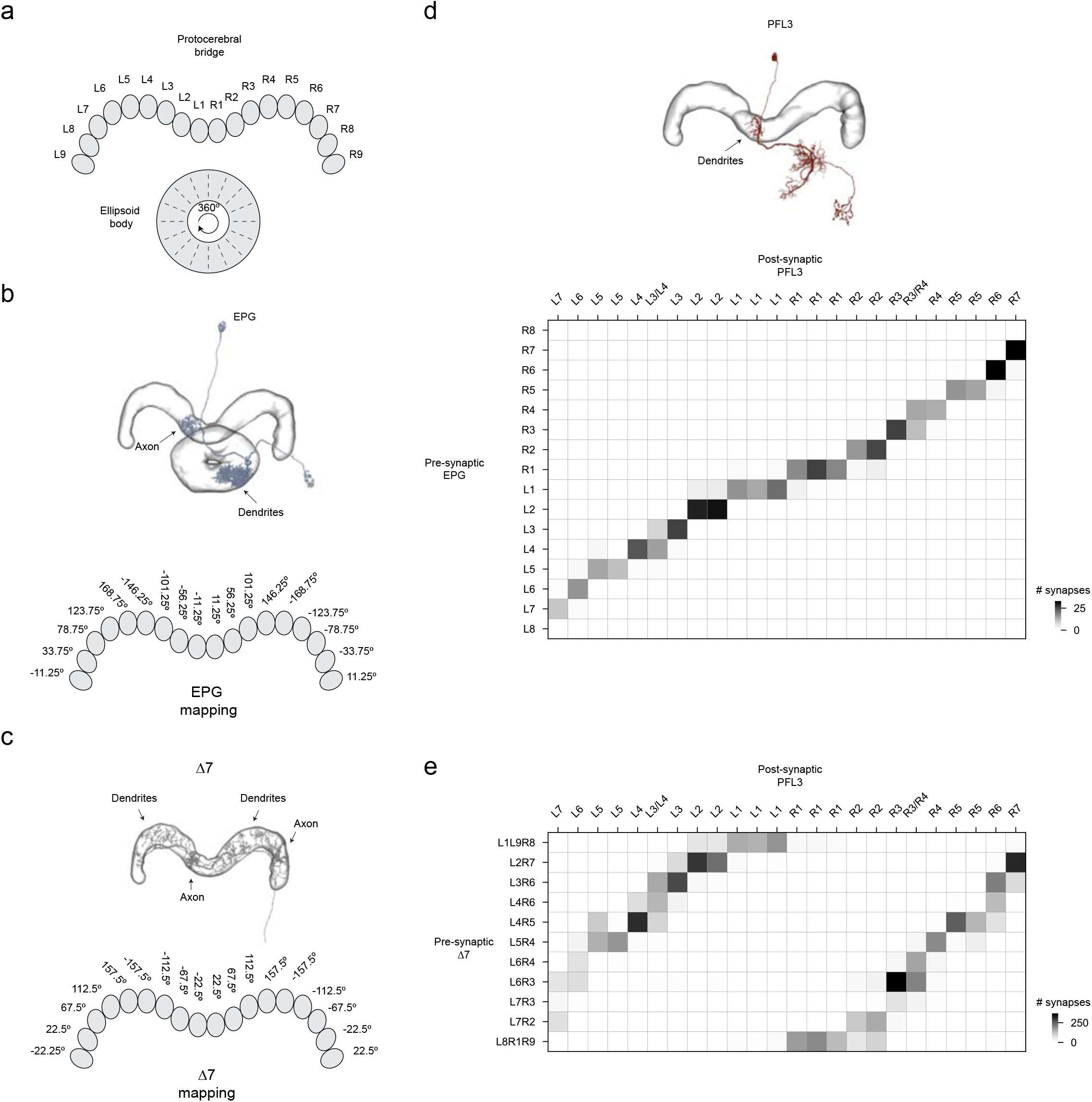
EPG and Δ7 connectivity with PFL3 neurons in the protocerebral bridge. PFL3 neurons receive inputs from two sets of heading-sensitive neurons in the protocerebral bridge: EPG neurons and Δ7 neurons. All data in this figure were extracted from the hemibrain connectome, neuPrint v1.2.1^15^. **a**, The ellipsoid body is divided into 16 wedges and the protocerebral bridge is divided into 18 glomeruli^34^. **b**, A single EPG neuron innervates one wedge of the ellipsoid body and projects to one glomerulus in the bridge (top). If one assumes that the ellipsoid body circle represents 360° of azimuthal space around the fly, consistent with physiological observations^2,37^, then each bridge glomerulus can be assigned an angle based on the wedge in the ellipsoid body from which the EPG cells that innervate that glomerulus originate (bottom) (see Lyu et al. 2022 for more details). The angles thus assigned to the bridge yield 45° azimuthal spacing between bridge glomeruli, except the inner two inner glomeruli, which are separated by only 22.5°. **c**, A single Δ7 neuron receives dendritic inputs (thin neurites in image) from EPG neurons across multiple glomeruli in the protocerebral bridge (see also Lyu et al. 2022 and Hulse et al. 2021) and expresses axonal terminals in 2-3 bridge glomeruli, spaced eight glomeruli apart. Two axon terminals are visible in the example Δ7 cell shown. If one counts the EPG->Δ7 synapses in each bridge glomerulus, and assume all EPG synapses are weighted the same, one can generate a predicted angle in which the output of each Δ7 cell should be maximal^4^, which is consistent with physiological observations^4,37^. These angles, based on the fact that individual Δ7 axons are offset from their dendrites by ∼180°, should yield angular assignments to bridge glomeruli that are ∼180° offset from the EPG assignments. However, because Δ7 cells are glutamatergic and appear to act in sign-inverting/inhibitory fashion on most of their downstream targets, their influence is actually expected to be roughly aligned with that of EPG cells, with a slight offset. Specifically, the Δ7 angular assignments in the bridge have a consistent 45° spacing between all glomeruli in the bridge, with a +ll.25° and −ll.25° offset relative to EPG angles for the right and left bridge respectively. **d**, EPG to PFL3 synaptic connectivity matrix. Each row represents a different group of 3-4, presynaptic, EPG cells, defined by their joint innervation in a given bridge glomerulus. Each column represents an individual postsynaptic PFL3 neuron, with twenty-four PFL3 cells in total. Canonical EPG neurons do not innervate the outer glomeruli of each side of the bridge and PFL3 neurons do not innervate the outer two bridge glomeruli on each side. The heatmap depicts the total number of synapses between each EPG-PFL3 pair. Note that PFL3 L3/L4 receives inputs from EPG neurons innervating both glomeruli L3 and L4. Likewise, PFL3 R3/R4 receives inputs from R3 and R4 EPG neurons. e, Δ7 to PFL3 connectivity matrix. Same as panel d, except that the rows depict different groups of Δ7 neurons rather than EPG neurons.

**Extended Data Fig. 7.**
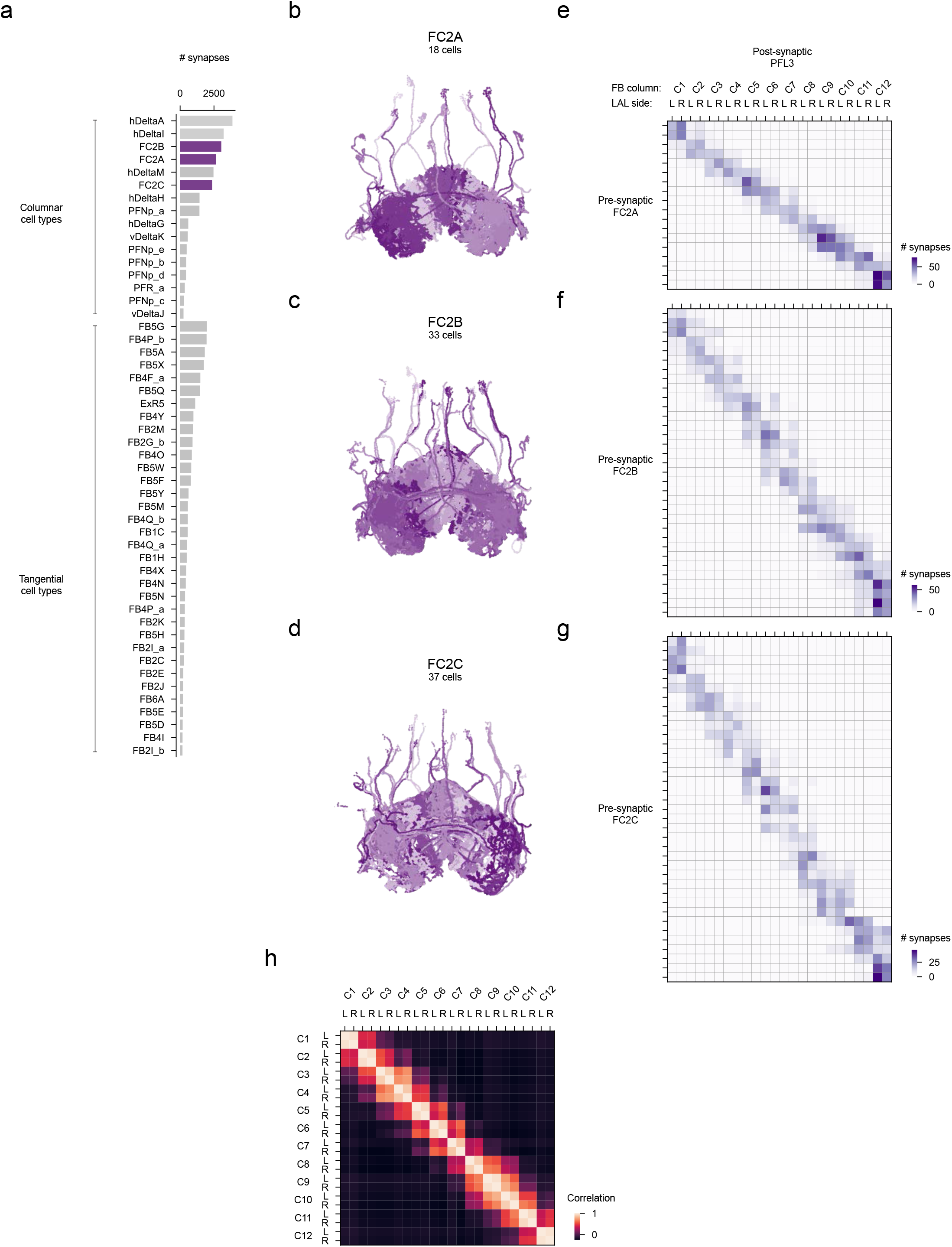
FC2 neurons represent a columnar-neuron class with a large number of synaptic inputs to PFL3 neurons. All data in this figure were extracted from the hemibrain connectome, neuPrint v1.2.1^15^. **a**, Plotted are the top 50 synaptic inputs to PFL3 neurons in the fan-shaped body. These cell classes constitute 94% of all PFL3 inputs in the fan-shaped body. Each row shows the total number of synapses between a pre-synaptic cell type and PFL3 neurons. FC2 neurons (purple) are a population of columnar neurons composed of three subtypes: FC2A, FC2B and FC2C. Together they constitute a third of columnar-cell synapses onto PFL3 cells in the fan-shaped body. Other columnar cell classes, such hDeltaA, hDeltaI, and hDeltaM cells could also provide goal information to PFL3 neurons during menotaxis or other goal-directed behaviours. Unlike columnar neurons, tangential cells have neurites that cut across all the columns of the fan-shaped body. These cells are likely to serve a role in modulating and impacting columnar goal information to the PFL3 cells, but their anatomy makes it less likely that they communicate column-specific information independent of their interaction with columnar neurons. **b-d**, Skeletons of FC2A, FC2B and FC2C populations. **e**, FC2A to PFL3 connectivity matrix. Each column represents an individual PFL3 neuron, sorted by its column in the fan-shaped body (C1 to C12) and whether it innervates the left (L) or right (R) LAL. C1 is on the very left of the fan-shaped body and C12 on the very right. Each row represents an individual FC2A neuron. **f**, FC2B to PFL3 connectivity matrix. **g**, FC2C to PFL3 connectivity matrix. h, Pairwise Pearson correlation matrix between individual PFL3 neurons based on their FC2 neuron inputs. The synaptic connections from all FC2 neurons to a given PFL3 neuron are treated as a vector and the correlation between each vector is computed. This analysis highlights that left and right PFL3 neurons innervating the same column receive highly similar inputs (which can also be seen by inspecting panels e-g). PFL3 neurons can be viewed as forming nine functional columns instead of twelve14. In this view, the four PFL3 neurons innervating the anatomical columns C3 and C4 (in the 12-column numbering scheme) would form a single functional column. The same would be true for C6 and C7, and C9 and C10. One justification for this scheme is that the four PFL3 neurons which would be combined to form a single functional column all share identical (or very similar) preferred-heading directions (see Fig. 4b). However, given that PFL3 neurons innervating C6 and C7, for example, receive different FC2 inputs, physiological evidence demonstrating that these FC2 inputs are in fact functionally identical would be required, we believe, to justify merging two anatomical columns into a single functional column and employing a 9-column fan-shaped body functional scheme instead of the 12-column scheme used herein.

**Extended Data Fig. 8.**
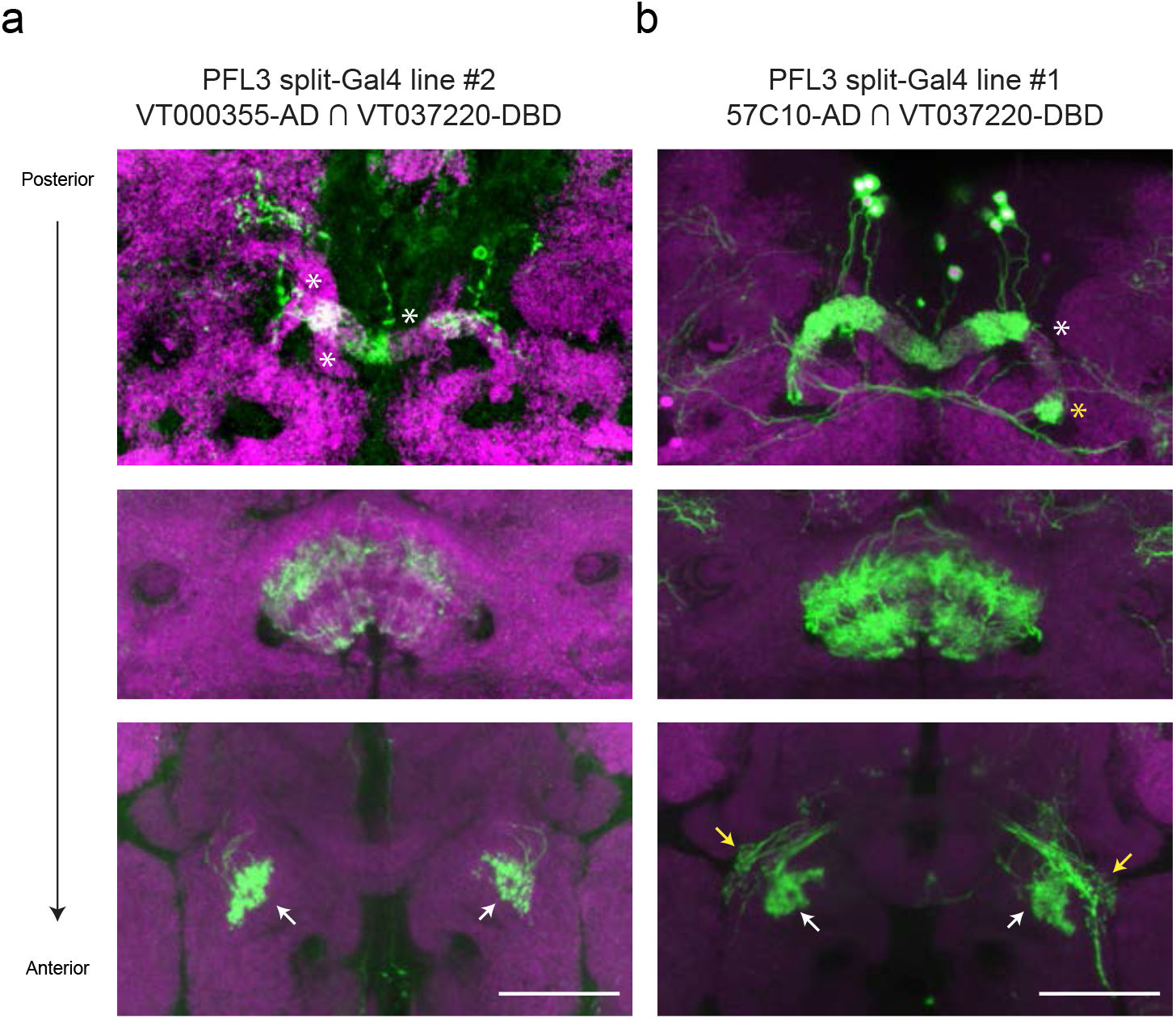
PFL3 split-Gal4 lines characterization. **a**, tdTomato expression (shown in green) driven in the VT000355-AD∩VT037220-DBD split-Gal4 line (used for patch-clamp and LAL-stimulation experiments) and anti-nc82 neuropil stain (magenta). Each panel shows an average z-projection. Top: protocerebral bridge. White asterisks highlight some of the glomeruli that are expected to have PFL3 innervation, but in which we could not detect fluorescence signal, indicating that the line does not label all PFL3 cells. Middle: fan-shaped body. Bottom: lateral accessory lobes. This line also stochastically labels PEG neurons. This was not a concern for either our patch-clamp (see Extended Data Fig. 9) or our LAL-stimulation experiments, since PEG neurons do not innervate the LAL. Scale bar is 50 μm. **b**, GFP expression driven in the 57C10-AD∩VT037220-DBD split-Gal4 line (used for LAL imaging experiments). Top: protocerebral bridge. The white asterisk highlights a glomerulus lacking clear PFL3 signal, indicating that the line does not label all 24 PFL3 cells. The yellow asterisk shows a glomerulus innervated by a non-PFL3 neuron (likely a PEG neuron), since PFL3 neurons do not innervate the outer two glomeruli. Middle: fan-shaped body. Bottom: lateral accessory lobes. White arrows highlight PFL3 expression in the left and right LAL. Yellow arrows mark non-PFL3 expression, which we excluded from our regions of interest for imaging analysis. Scale bar is 50 μm.

**Extended Data Fig. 9.**
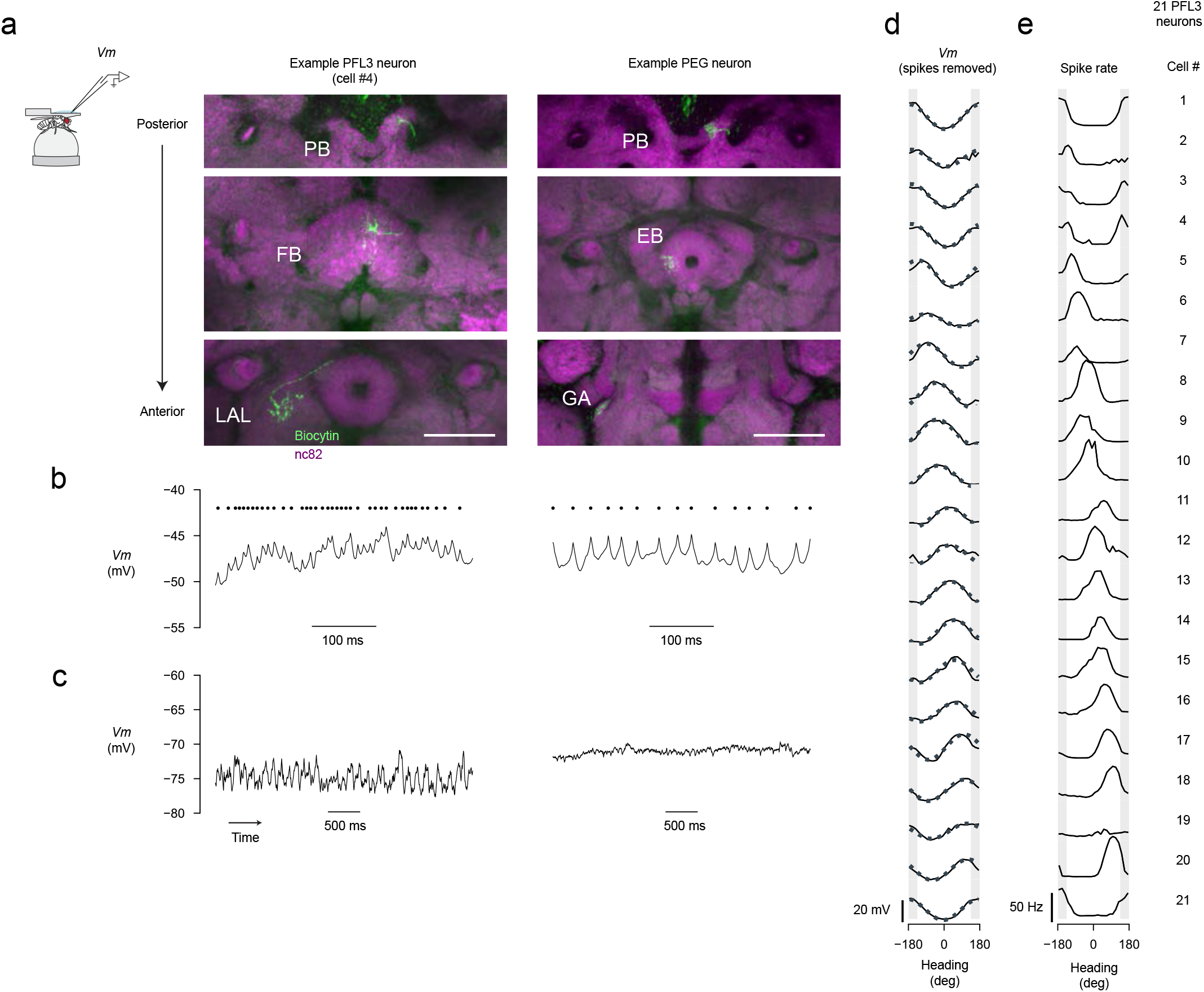
PFL3 neurons can be distinguished from PEG neurons based on their electrophysiological properties and individual PFL3 neurons are tuned to heading, with different cells showing different preferred heading angles. **a**, Biocytin fill of a PFL3 neuron (left) and a PEG neuron (right) recorded in the split-Gal4 line VT000355-AD∩VT037220-DBD. PEG and PFL3 neurons can be differentiated based on their innervation patterns. Specifically, PFL3 neurons innervate the fan-shaped body (FB) and lateral accessory lobe (LAL) whereas PEG neurons innervate the ellipsoid body (EB) and the gall (GA). Each image is an average z-projection from a subset of slices. Scale bar is 50 μm. **b**, Sample *Vm* from the PFL3 and EPG neuron depicted in the anatomy panels directly above. At depolarized membrane potentials, the spikes of PFL3 neurons were relatively small (left) whereas those from the PEG neurons were relatively large (right). Black dots indicate detected spikes. **c**, At hyperpolarized membrane potentials, PFL3 neurons display rhythmic oscillations (left), whereas the membrane potential of PEG neurons tends to be more flat (right). **d-e**, *Vm* (spikes removed) (left) and spike rate (right) tuning curves to heading direction for all PFL3 cells. Dashed line in the *Vm* curves represents a sinusoidal fit to data, which was used for estimating the cell’s preferred-heading direction (see Methods). Shaded area represents 90º gap at the back of the arena where the bar is not visible. Cells are sorted and numbered based on their estimated preferred-heading direction. We use this numbering scheme throughout the manuscript to refer to specific cells.

**Extended Data Fig. 10.**
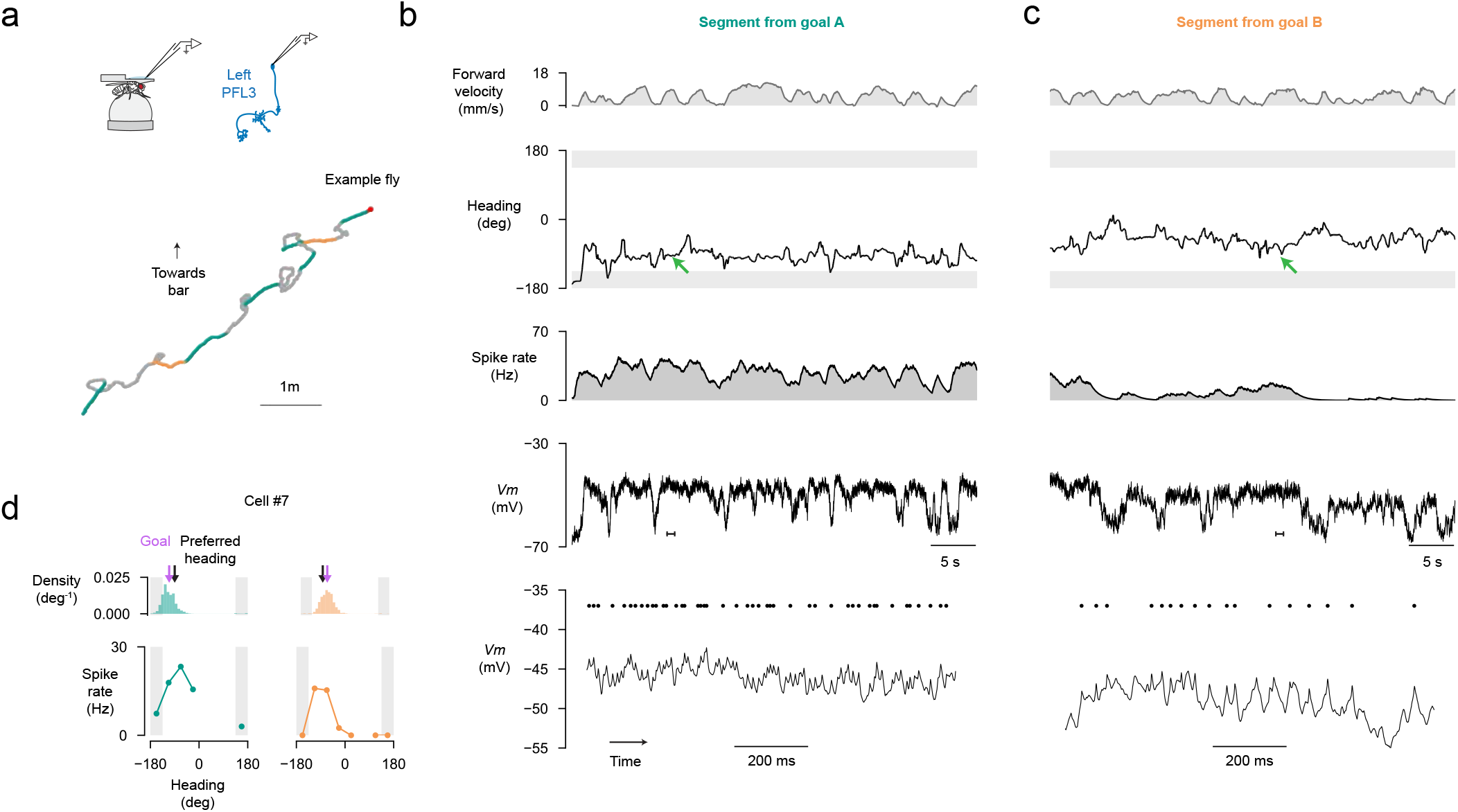
An example PFL3 neuron shows goal-dependent scaling of its tuning curve to heading. **a**, Trajectory of a fly from which we recorded a PFL3 neuron that projected to the left LAL. Colored portions of the trajectory correspond to menotaxis bouts where the fly’s goal angle relative to the cell’s preferred-heading direction was between −45º and 0º (turquoise) or between 0º and 45º (orange), see Methods for details. Red dot marks the start of the fly’s trajectory. **b**, Top row, example timeseries from when the fly’s goal angle was to the left of the cell’s preferred-heading direction (corresponding to a portion of the turquoise traces). Second row, heading over time. Green arrows mark a moment when the fly experienced a similar heading direction for this goal angle and the goal angle in the next panel. Bottom row: expanded view of membrane potential corresponding to the period marked by the inset in the third row. Spikes are marked with black dots. Note that the spike rate is higher in this panel, when the fly’s goal was to the left of the cell’s preferred direction compared to the spike rate in the next panel, despite the fly experiencing similar heading directions in the two situations. **c**, Same as panel **b**, except this is an example timeseries from when the fly’s goal was to the right of the cell’s preferred direction (corresponding to a portion of the orange traces). **d**, Top: Histogram of the fly’s heading for the menotaxis bouts highlighted in panel **a**. Bottom: spike rate tuning curves to heading for the menotaxis bouts whose behavioral heading distributions are indicated in the histograms above.

**Extended Data Fig. 11.**
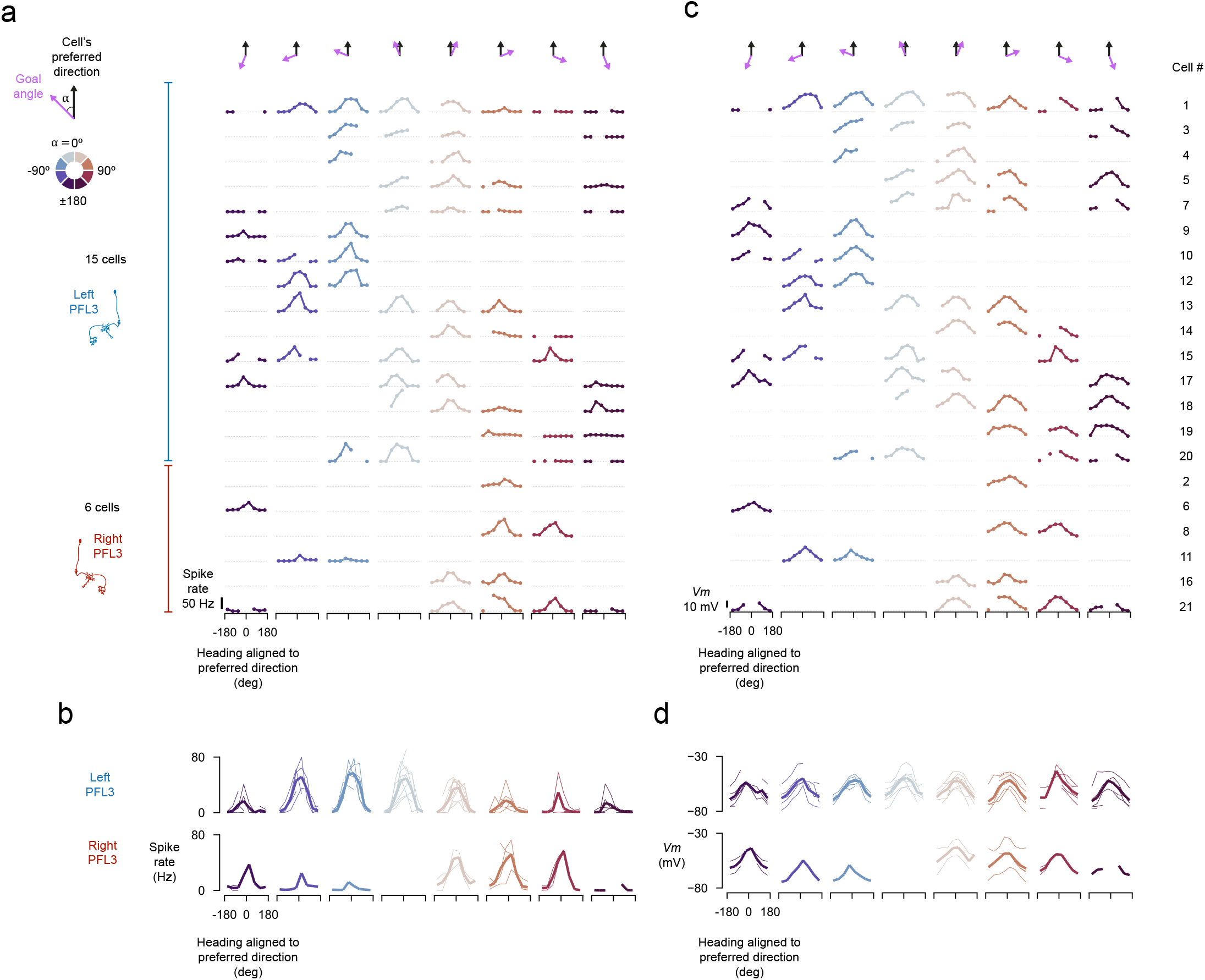
Goal-dependent scaling of PFL3 activity is more prominent in the spike rate than in the somatic membrane potential. **a**, After determining a cell’s preferred heading angle from the overall tuning curve (Extended Data Fig. 9d), we plotted a set of tuning curves with a shifted *x*-axis for each cell, so as to always have this preferred angle at zero. Here we show such preferred-phase nulled tuning curves binned by the fly’s goal angle relative to the cell’s preferred direction. Each row represents a different cell. Each column (and color) represents a different bin of goal angles relative to cell’s preferred direction, with the mean angle of that bin represented by the purple arrow. Because single flies typically adopted only a few goal directions throughout a recording session, this led to the many “missing” tuning curves seen here. Likewise, some tuning curves are missing data in some portions of the *x*-axis because for each goal direction, a fly does not typically experience the full range of heading directions, even with our bar jumps aiming to minimize this issue. Dashed grey line indicates a spike rate of zero Hz. **b**, Spike rate mean across all cells. Thin lines: individual cells. Thick line: mean across cells. Top row is the same as Figure. 1k. **c**, Same as in panel **a** but plotting membrane potential (spikes removed) (Methods). For each row (i.e. cell), the grey dotted line represents the row’s minimum membrane potential. The “cell #” identifiers shown on the right are identical to those used in Extended Data Fig. 9 and these numbers apply also to panel **a. d**, Mean membrane potential (spikes removed) across all cells. These plots were generated by averaging the raw membrane potential, which was corrected for the same 13 mV liquid-liquid junction potential across all recordings, but not shifted by the minimum membrane potential for each cell prior to averaging. Thin lines: individual cells. Thick line: mean across cells.

**Extended Data Fig. 12.**
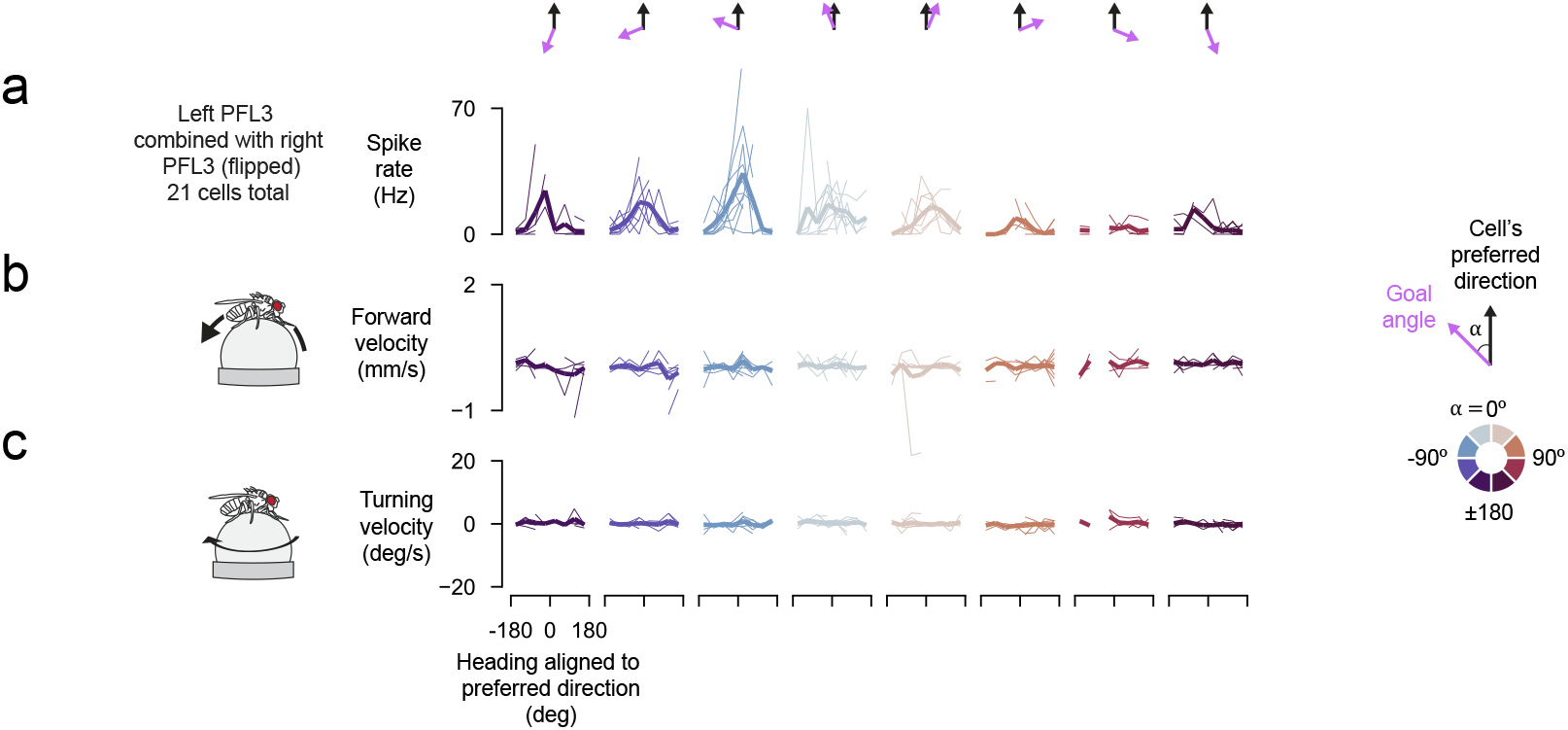
The goal-dependent scaling of PFL3 spike-rate tuning curves is not a simple consequence of the fly’s instantaneous walking dynamics. **a**, The PFL3 spiking activity and the fly’s locomotor behaviour were analyzed by binning the data according to the fly’s goal relative to the cell’s preferred direction (columns) and the fly’s heading angle relative to the cell’s preferred direction (*x*-axis of each tuning curve) as in Extended Data Fig.11. The difference here is that we only included timepoints when the filtered forward walking velocity of the fly was below 0.5 mm/s and the fly’s turning velocity was between −5 °/s and 5 °/s. We lumped our six recordings from right PFL3 neurons with our 16 recordings from left PFL3 neurons, by flipping the heading-relative-to-the-cell’s-preferred-heading and goal-heading-relative-to-the-cells’-preferred-heading values for right PFL3 neurons prior to averaging. Thin lines: individual cells; thick line: mean across cell. **b**, Mean forward walking velocity, analyzed as described in panel **a. c**, Mean turning velocity, analyzed as described in panel a.

**Extended Data Fig. 13.**
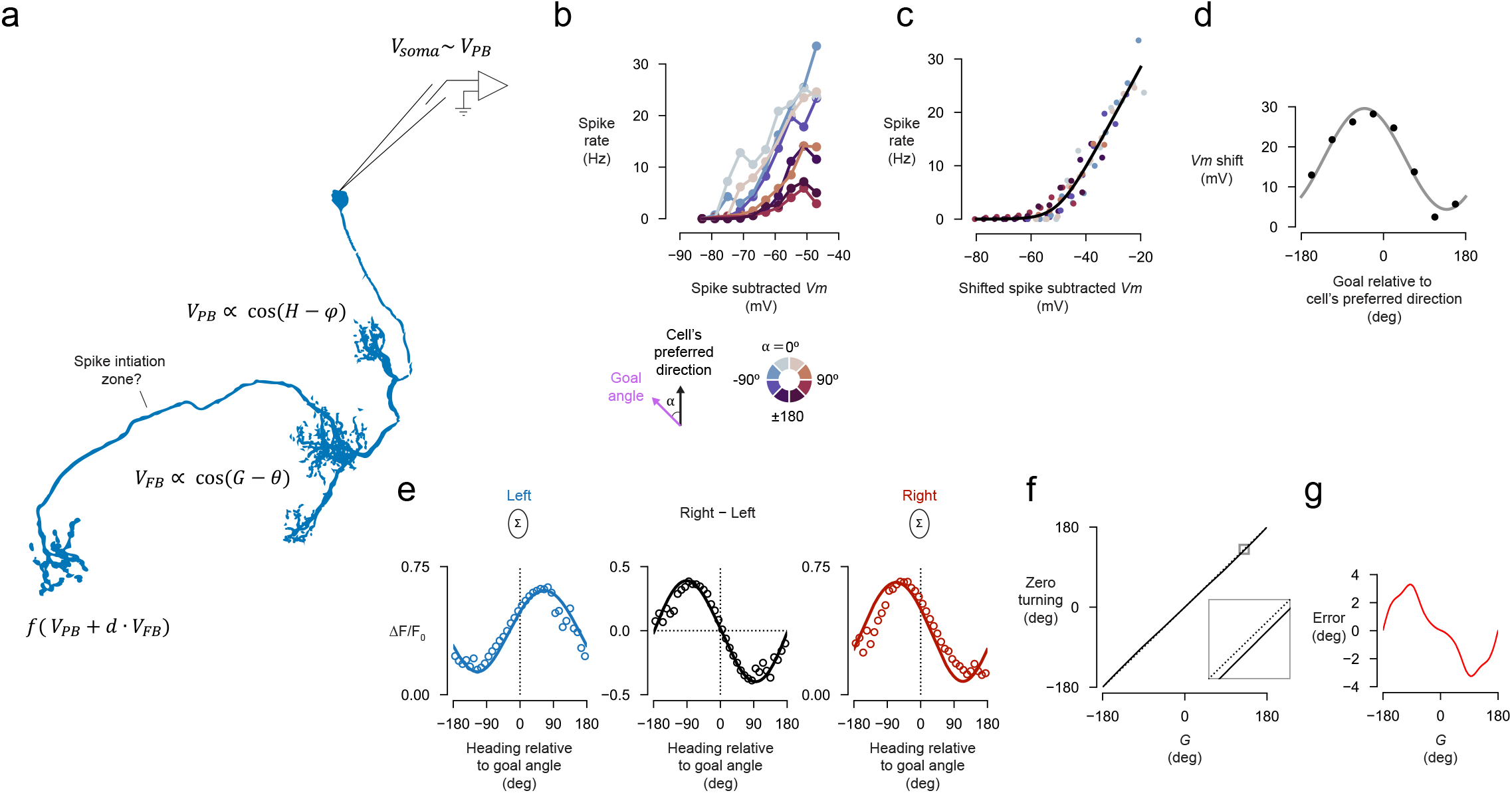
Model for how heading and goal information is integrated in individual PFL3 neurons. **a**, Schematic for how PFL3 neurons integrate heading and goal information. Two inputs contribute to the membrane potential of a PFL3 cell. One input comes from the protocerebral bridge and yields a membrane potential signal, VPB, in the PFL3 cell that is proportional to a cosine function of the difference between the fly’s heading angle, *H*, and the PFL3 cells’ preferred heading angle, φ. The other input comes from the fan-shaped body and results in the membrane potential signal, *V*_*FB*_, in the PFL3 cell that is a cosine function of the different between the fly’s goal angle, G, and the cell’s preferred goal angle, θ. The membrane potential measured at the soma, *Vm*, is dominated by VPB because the fan-shaped body is electrotonically further away from the soma than the protocerebral bridge (consistent with the more modest goal-dependent changes *Vm* we observed in Extended Data Fig. 11). The spike rate of the neuron is given by a nonlinear function of a sum of the cosine functions describing *V*_*PB*_ and *V*_*FB*_ (with *V*_*FB*_ scaled by a weighting factor *d*, refecting the relative strengths of these two inputs at the spike initiation zone). **b**, Spike-rate vs *Vm* (spikes removed) for different goal angles relative to the cell’s preferred heading from our whole-cell recordings. The fact that this relationship depends on the fly’s goal angle, is, we believe, due the somatic membrane potential predominantly reflecting heading input from the bridge; we assume in the model that the spike-initiation zone has access to both the heading- and goal-related inputs. Each dot shows the mean across cells (right PFL3 neurons were included by flipping the sign of the goal-to-preferred heading angle, see Methods). **c**, The same curves as in panel b, but shifted along the horizontal axis in order to maximally align them (Methods). The black curve is a softplus function fit to the data points (Methods). **d**, The shifts from panel c, plotted as a function of the goal angle of the corresponding spike-rate curve. The fact that these shifts have a cosine shape as a function of the goal angle is consistent with: (1) the existence of a cosine-shaped goal input in the fan-shaped body (as our model assumes) and (2), our hypothesis that the voltage consequences of the goal in the fan-shaped body are not fully evident in the soma, thus requiring the *Vm* shift in the plot in panel b, to align all the curves to a common spike-rate vs. *Vm* underling function, as is assumed in our model. **e**, A direct comparison of model predictions (Fig. 4f) and calcium imaging results (Fig. 5D) for right and left LAL signals and for the turning signal. **f**, The black curve shows the predicted relationship between the ‘zero’ heading (i.e. the heading angle where the turning signal is zero and the slope is negative) and *G* (the phase of the goal signal). The inset shows an expanded view of the area marked by the small grey box. Note that zero heading and *G* are not exactly equal (dotted line is the diagonal). **g**, The angular difference between the zero heading and *G* (error) as a function of G.

**Extended Data Fig. 14.**
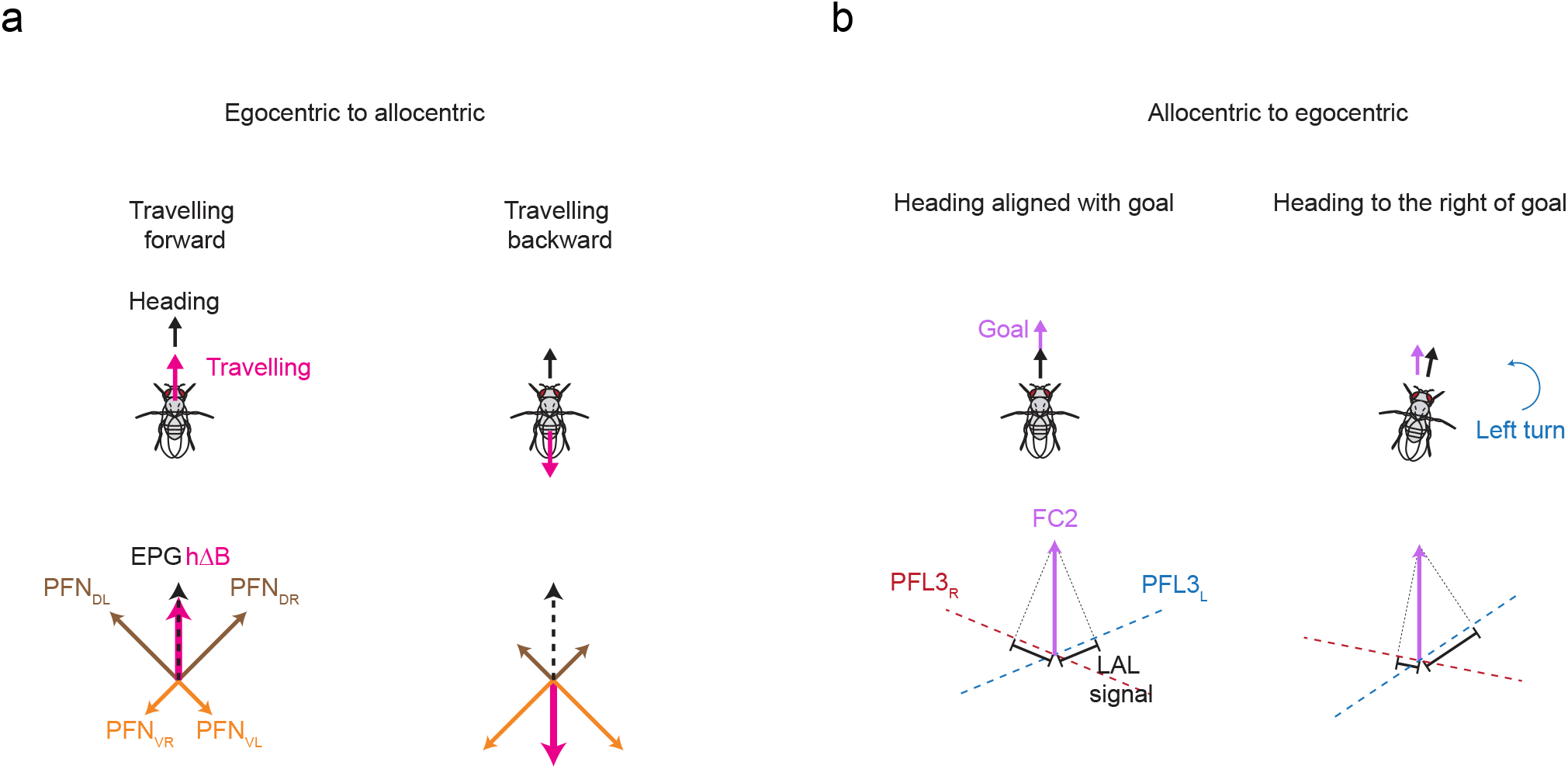
Schematic models for how the fly’s brain performs egocentric-to-allocentric and allocentric-to-egocentric coordinate transformations. **a**, The PFNd/PFNv circuit converts the fly’s egocentric traveling direction, as signaled in sensory inputs to the central complex, into an allocentric-traveling direction signal in hB cells (adapted from Lyu et al. 2022). Two arrays of PFNd cells and two arrays of PFNv cells express sinusoidal activity patterns whose phase and amplitude can be considered as representing four vectors with a specific angle and length (brown and orange vectors). To calculate the allocentric traveling direction, the four neurally represented vectors all initially signal the angle in which the fly is translating in relation to its body axis, with the amplitude of each activity pattern representing a projection of that egocentric traveling vector onto a different (basis) direction. All four vectors are then rotated together based on the fly’s heading relative to external cues, (e.g., the sun), which implements the egocentric-to-allocentric transformation. Finally, the circuit finds the max position of the summed, output vector, which represents the fly’s traveling angle in reference to external cues. When the fly is traveling forward, the two forward-facing PFNd vectors are long and the two backward-facing PFNv vectors are short, yielding an output traveling vector in hB cells that points in the direction of the fly’s heading, as encoded by the EPG bump (left schematic). When the fly is traveling backward, the two, forward-facing, PFNd vectors are short and the two, backward-facing, PFNv vectors are long, yielding an output traveling vector in hB cells that points in a direction 180° opposite of the fly’s heading, as encoded by the EPG bump (right schematic). **b**, Left: The PFL3 circuit that converts an allocentric goal angle into an egocentric steering signal can be considered, computationally, to be taking the difference between two dot products. The left and right PFL3 neurons form two non-orthogonal axes (blue and red dotted lines). Each axis represents the fly’s heading angle rotated either clockwise and counter-clockwise by the same angle. The fly’s allocentric goal angle, signaled by the position of the FC2 bump in the fan-shaped body, is represented by the purple vector. The projection of the goal vector onto the blue PFL3 axis (which can be considered as the output of a dot product between the goal vector and unit vector pointing along the blue axis) reflects the sum of the right PFL3 activity in the LAL (and vice versa for the left PFL3 axis). When the fly is aligned with its goal, the difference between the red and blue dot products is zero. Right: When the fly changes its heading, the axes rotate and the difference between the two now tells the fly to turn left. Neuronally, the left and right PFL3 axes represent vectors generated by projecting their heading inputs in the bridge onto the fan-shaped body. The amount by which the left and right PFL3 axes are rotated is determined by the anatomical shift in the PFL3 projection pattern from the bridge to the fan-shaped body.

## Methods

### Fly husbandry

*Drosophila melanogaster* flies were raised at 25°C on a 12-hour light/dark cycle. All physiological experiments were performed on 1 to 3 day old female flies. For optogenetic experiments, experimental and control crosses were kept in a box with a blue gel filter (Tokyo Blue, Rosco) as a cover—to minimize exposure to light within the excitation spectrum of CsChrimson while also not keeping the flies in complete darkness; eclosed flies from such experiments were placed onto food containing 400 μM all-trans retinal for at least one day.

### Fly genotypes

To image EPG neurons during mentoaxis experiments (Fig. 1, Extended Data Fig. 3), we used +/-; 60D05-Gal4/+; UAS-GCaMP7f/+.

To image FC2 neurons during menotaxis experiments (Fig. 1, Extended Data Fig. 3) we used either +; VT065306-AD/+; VT029306-DBD/UAS-GCaMP7f or +; VT065306-AD/+; VT029306-DBD/UAS-sytGCaMP7f.

To stimulate FC2 neurons while imaging (Fig. 2, Extended Data Fig. 5) we used +; VT065306-AD/UAS-CsChrimson-tdTomato; VT029306-DBD/UAS-sytGCaMP7f. For control flies we used +; VT065306-AD/UAS-Tomato; VT029306-DBD/UAS-sytGCaMP7f.

To label PFL3 neurons for patch-clamp experiments (Fig. 3, Extended Data Figs.9-13) we used: +; VT000355-AD/UAS-2xeGFP; VT037220-DBD/+.

To label PFL3 neurons for calcium imaging only (Fig. 5a-d) we used +; 57C10-AD/UAS-Tomato; VT037220-DBD/UAS-GCaMP7f.

To stimulate PFL3 neurons while imaging (Fig. 5e-h) we used +; VT000355-AD/UAS-GCaMP7f; VT037220-DBD/UAS-CsChrimson-tdTomato. For control flies we used +; VT000355-AD/UAS-Tomato; VT037220-DBD/UAS-GCaMP7f (Fig. 5g-h).

To stimulate PFL1 neurons while imaging (Fig. 5g-h) we used +; VT000454-AD/ UAS-GCaMP7f; VT001980-GAL4/ UAS-CsChrimson-tdTomato.

To characterize the expression pattern of VT065306-AD; VT029306-DBD (Extended Data Fig. 1a) and 57C10-AD; VT037220-DBD (Extended Data Fig. 8b) we crossed each of these lines to UAS-UAS-RedStinger; UAS-mCD8-GFP.

To characterize the expression pattern of VT000355-AD; VT037220-DBD we crossed this line to UAS-Tomato (Extended Data Fig. 8a).

For multicolor flip-out of VT065306-AD; VT029306-DBD we used hs-FLPG5.PEST (Extended Data Fig. 1b).

### Origins of fly stocks

We obtained the following stocks from the Bloomington Drosophila Stock Center, the Janelia FlyLight Split-Gal4 Driver Collection or from other labs:

VT000454-p65AD; VT001980-GAL4.DBD (SS02239)^47^

VT000355-p65AD (attP40)^47^

57C10-p65AD (attP40) (BDSC #70746)

VT037220-Gal4.DBD (attP2) (BDSC #72714)

R60D05-Gal4 (attP2) (BDSC #39247)

UAS-2xeGFP (Dickinson lab)

20XUAS-IVS-jGCaMP7f (VK05) (BDSC #79031)

10XUAS-sytGCaMP7f (attP2) (BDSC #94619)

UAS-CsChrimson-tdTomato (VK22) (gift from David Anderson, Barret Pfeiffer, and Gerry Rubin)

UAS-CsChrimson-tdTomato (VK05) (gift from David Anderson, Barret Pfeiffer, and Gerry Rubin)

UAS-mCD8-GFP (attP2) (BDSC # 32194)

UAS-RedStinger (attP40) (BDSC #8546)

hs-FLPG5.PEST (BDSC #64085)

### Generation of genetic driver lines and immunohistochemistry

To generate split-Gal4 lines targeting FC2 and PFL3 neurons, we used both the Color MIP tool^48^ and NeuronBridge^49^ to find suitable pairs of hemi-driver lines. We validated that the split-Gal4 lines generated target the cells of interest by means of immunohistochemistry (Extended Data Fig. 1 for FC2 cells, Extended Data Fig. 8 for PFL3 cells).

We dissected and the brains incubated them in either 2% paraformaldehyde (PFA) for 55 min. at room temperature or in 1% PFA overnight at 4°C. We blocked and de-gassed brains in a blocking solution consisting of 5% normal goal serum (NGS) in 0.5% Triton X-100, phosphate buffered saline (PBT).

For GFP or tdTomato labeling experiments (Extended Data Figs. 1a,8), we used a primary antibody solution of 1:100 chicken anti-GFP, 1:500 anti-dsRed and 1:10 mouse anti-nc82 in 1% NGS-PBT and a secondary antibody solution consisting of 1:800 goat anti-chicken:Alexa Fluor 488, 1:400 anti-rabbit 594 and 1:400 goat anti-mouse:Alex Fluor 633 in 1% NGS-PBT.

For heat-shock multicolor flip-out experiments^50^ (Extended Data Fig. 1b), we used a primary antibody solution of 1:300 rabbit anti-HA, 1:200 rat anti-FLAG and 1:10 mouse anti-nc82 in 1% NGS-PBT. The secondary antibody solution used was 1:500 donkey anti-rabbit:Alexa 594, 1:500 donkey anti-rat:Alexa 647 and 1:400 goat anti-mouse:Alexa Fluor 488 in 1% NGS-PBT, followed by a tertiary antibody solution of 1:500 DyLight anti-V5 549 in 1% normal mouse serum PBT.

For visualizing biocytin-labeled neurons after patch-clamp experiments (Extended Data Fig. 5a), the primary antibody solution we used was 1:10 mouse anti-nc82 in 1% NGS-PBT and the secondary antibody solution was 1:800 goat anti-mouse:Alex Fluor 488 and 1:1000 streptavidin:Alexa Fluor 568 in 1% NGS-PBT.

Brains were mounted in Vectashield and images were acquired using a Zeiss LSM780 confocal microscope with 40x 1.20 NA objective.

### Fly tethering and preparation

We glued flies to custom holders that allowed for physiological measurements from the brain, under a saline bath, while the body remained dry and capable of executing tethered locomotor behaviour, as described previously^37,38^. When imaging neuronal activity in the protocerebral bridge or performing electrophysiology, we tilted the fly’s head down such that the brain was viewed from the posterior side. When imaging neuronal activity in the lateral accessory lobes or the fan-shaped body, the fly’s head was not tilted and the brain was viewed from the dorsal side. Glue was added at the junction of the fly’s thorax and wings to prevent tethered flight and, the proboscis was glued to the head to minimize brain motion associated with large proboscis movements. Brains were exposed by cutting and removing a small piece of cuticle with a 30-gauge syringe needle followed by removal of trachea and fat cells overlying the brain with forceps.

A previous study noted that wild-type flies typically perform menotaxis behaviour when food deprived for 8-16 hours and heated to 34°C^13^. In the present study, we noticed that for some genotypes, the same level of food deprivation would yield unhealthy flies. As such, we opted for a shorter period of food deprivation for most experiments. We typically performed experiments at least three hours after tethering flies. During this interval, we kept tethered flies inside a box with a wet piece of tissue paper to prevent desiccation. For FC2 stimulation experiments, we placed flies on plain agarose roughly 14 hours before tethering. In all experiments, we heated the tethered fly by perfusing 26-30°C saline over the fly’s head using a closed-loop temperature control system (Warner Instruments, CL-100).

### Virtual reality setup

For both two-photon calcium imaging and patch-clamp experiments, we placed flies in a virtual reality setup described previously^38^. In brief, tethered flies were positioned over an air-cushioned foam ball^2,38^ (Last-A-Foam FR-4618, General Plastics) that had a diameter of 8 mm. The ball’s movements were visualized with a Chameleon CM3-U3-13Y3M (Teledyne FLIR) camera, whose 3D pose was tracked at 50 Hz using FicTrac^51^. We used a cylindrical LED display that spanned 270° of angular space around the fly^39^. In all experiments, the fly’s yaw rotations on the ball controlled the position of an 11°-wide vertical blue bar^38^.) We covered the arena with sheets of blue gel filter (Tokyo Blue, Rosco) in order to prevent blue light bleed-through into the photomultiplier tubes. In patch-clamp experiments, we placed a steel mesh in front of the arena to electrically shield the headstage, as well as a nylon mesh to minimize reflections.

### Calcium imaging

We performed two-photon calcium imaging as described previously^38^, with certain changes indicated below. We used a Scientifica Hyperscope and a Chameleon Ultra II Ti:Sapphire femtosecond pulsed laser (Coherent) tuned to 925 nm. We performed volumetric imaging, using galvo-galvo mode (Cambridge Technologies MicroMax) to scan the *xy*-plane and a piezo device (PI, P-725.4CA) to move a 16x/0.8 NA Nikon objective along the *z*-axis. Emission light was split using a 565 nm dichroic mirror. We used a 500-550 nm bandpass filter for the green signal and a 590-650 nm bandpass filter for the red signal. Emission photons were detected and amplified using GaAsP detectors (Hamamatsu, H10770PA-40). ScanImage^52^ (2018b) software was used to control the microscope.

For Fig. 5a-d, we used ScanImage’s MultipleROI feature to define two 50 × 50 pixel regions of interest (ROI) for each side of the lateral accessory lobes (LAL). We scanned the LAL with two *z* slices per volume, yielding a volume rate of 9.16 Hz. For Fig. 1 we scanned the protocerebral bridge or the fan-shaped body at 4.95 Hz using a 128 × 64 pixel ROI with 3 *z* slices. In standard imaging experiments (Fig. 1 and Fig. 5a-d), we used a laser power of ∼25 mW (measured after the objective). Imaging experiments lasted up to 26 minutes. Occasionally, the fly’s brain would slowly sink over the course of a recording. To correct for this motion, we manually adjusted the position of the objective via a microscope-stage motor during the recording.

### Optogenetic stimulation during imaging

We used the same two-photon light path to image and focally stimulate neurons, using ScanImage’s MultipleROI feature. We defined two ROIs which we refer to as the imaging ROI and the stimulation ROI (Extended Data Fig. 5a). The imaging ROI included the entire structure of interest (lateral accessory lobe or fan-shaped body). We scanned this ROI with a low laser power (10 mW), which did not change throughout the recording. The stimulation ROI was smaller than the imaging ROI. We scanned the stimulation ROI with a higher laser power (50 or 70 mW) and the location of this ROI changed throughout a recording. Within each *z* slice, we first scanned the imaging ROI and then the stimulation ROI. We only used pixel values from the imaging ROI for the analysis of fluorescence changes. We used a MATLAB script to change the location of the stimulation ROI automatically during an experiment.

For Fig. 2, we alternated between stimulating one of two positions in the fan-shaped body (referred to as location A and B). We stimulated a more anterior position in the brain, which lacked CsChrimson-tdTomato expression, between trials (Extended Data Fig. 5b). This step ensured that the average laser power per volume remained constant throughout the experiment, which is important because flies could show behavioural reactions to changes in illumination intensity. To register the timing of a change in the location of the stimulation ROI, we recorded the *x* and *y* galvo positions over time. We used a stimulation power of ∼50 mW in these experiments. We imaged three *z*-slices and the stimulation ROI existed in all three slices. The acquisition rate was 3.32 Hz. The duty cycle was ∼0.67 (the number of pixels in the stimulation ROI divided by the total number of scanned pixels). If we acquired more than one recording per fly, the locations of the stimulation and imaging ROIs were adjusted as needed between recordings.

For Fig. 5e-h, we alternated from stimulating the left or right LAL. Between trials, we moved the stimulation ROI to a location anterior to the LAL that did not have any CsChrimson-tdTomato expression. We used a stimulation power of ∼70 mW in these experiments. We used a single *z*-slice to scan the LAL with an acquisition rate of 4.97 Hz and the duty cycle was ∼0.33.

We used a lower laser power in the imaging ROI so as to minimize two-photon excitation of CsChrimson. However, we noticed that during the inter-trial period the FC2 activity sometimes appeared non-physiological. For instance, the middle columns of the fan-shaped body, which are located more superficially, sometimes appeared to be persistently active during the inter-trial period, irrespective of the fly’s behaviour (e.g. Fig. 2c). We therefore suspect that at even low laser intensities we might have been optogenetically stimulating neurons to some extent. We therefore did not analyze the fly’s behaviour during inter-trial periods during these experiments, as these were associated with unphysiological activation of the system.

### Patch-clamp electrophysiology

We performed patch clamp experiments as described previously^37^, with some changes indicated below. We perfused the brain with an extracellular solution^53^ bubbled with carbogen (95% O_2_, 5% CO_2_). The composition of the extracellular solution (in mM) was: 103 NaCl, 3 KCl, 5 TES, 10 trehalose dihydrate, 10 glucose, 2 sucrose, 26 NaHCO_3_, 1 NaH_2_PO_4_, 1.5 CaCl_2_ and 4 MgCl_2_ (280±5 mOsm). The composition of the intracellular solution^53^ was (in mM): 140 K-Aspartate, 1 KCl, 10 HEPES, 1 EGTA, 0.5 Na_3_GTP, 4 MgATP (pH 7.3, 265 mOsm). For some recordings the solution also included 13 mM biocytin hydrazide and 20 mM Alexa-568, which could be used to fill the neuron for subsequent verification of the identity of the cell from which we were recording.

We illuminated the fly’s brain via an 850 nm LED (Thorlabs) coupled to an achromatic lens pair (MAP10100100-A, Thorlabs) that focused the LED’s light onto a small spot on the fly’s head. We used borosilicate patch pipettes (BF150-86-7.5, Sutter Instruments) with resistances of 6-13 MΩ. Recordings were conducted in current-clamp mode (MultiClamp 700B, Molecular Devices) with zero injected current. The voltage signal was low-pass filtered at 4 kHz before sampling at 10 kHz. Plots have been corrected for a 13-mV liquid-liquid junction potential. For recordings in which we included biocytin hydrazide and Alexa-568 in the intracellular solution, we visualized the recorded, filled cell, by taking a manual *z*-stack on our epi-fluorescence patch-clamp microscope while illuminating with a 565 nm LED (pE-100, CoolLED). We also dissected the brain and performed immunohistochemistry, staining for biocytin, to to verify the patched cell’s identity and anatomy.

Because the split-Gal4 line that we used for patch-clamp experiments (VT00355-AD ∩ VT037220-DBD) labels both PFL3 and PEG neurons (Extended Data Fig. 9a), we initially verified the cell type identity of all cells to be included in this paper via immunohistochemistry. Three PEG neurons and eight PFL3 neurons were identified by this method. Since recordings of verified PFL3 and PEG neurons were clearly distinguishable by their spike amplitudes and resting potential dynamics (Extended Data Fig. 9a-c), we classified the remaining recordings based on these electrophysiological criteria (7 PEG neurons and 13 PFL3 neurons).

To help categorize a recorded PFL3 neuron as innervating the left or right LAL, we targeted PFL3 cells with somas far from the midline as these PFL3 cells project exclusively to the contralateral LAL. Of the eight PFL3 neurons whose anatomy we verified via immunohistochemistry, all projected to the contralateral LAL. For an additional two PFL3 neurons we were able to verify that they projected contralaterally via the epifluorescence *z*-stack. We classified the remaining 11 PFL3 neurons based on their soma location. We discarded one recording from a soma located close to the midline since its identity as a left or right PFL3 could not be definitively established.

Because our recordings could approach two hours in length, we sometimes observed, a slow depolarizing drift in the membrane potential over time, accompanied by a decrease in spike size, consistent with a slowly increasing access resistance. We trimmed these recordings by visually inspection to only include the portion in which the membrane potential and spike size were stable. Four cells were discarded as there was no period when these criteria were met. After trimming, the average recording duration was 46 min (ranging from 6 min to 120 min).

### Experimental structure

In all experiments, we allowed the fly to walk in closed-loop with the bar for approximately 5-30 minutes as we prepared for data collection (i.e., during desheathing and seal attempts in patch-clamp measurements or during ROI selection in imaging experiments). This time period gave the fly experience with all possible angular bar positions, which is expected to reinforce the formation of a stable map between the position of the bar on the screen and the EPG heading-estimate in the central complex^54,55^. For experiments in which we did not employ optogenetics (Fig. 1, Fig. 3 and Fig. 5a-d), we used bar jumps to periodically assess whether the fly was actively maintaining its heading direction. Bar jumps served the additional role of ensuring that a fly sampled heading angles away from its goal angle, which allowed us to generate tuning curves to heading. Specifically, every 2 minutes, we instantaneously repositioned the bar by ±90° from its current position. The bar then remained static at this new location for 2 s, after which it returned to being under closed-loop control by the fly. For Fig. 1 and Fig. 5a-d each recording included five +90° bar-jump events and five –90° bar-jump events, presented in a random order. We typically collected two recording files from a given fly (a few flies had one or three recordings). In electrophysiology experiments, which could sometimes run as long as two hours, bar jump events occurred throughout, until the end of an experiment.

For the stimulation experiments in Fig. 2, each recording consisted of five location-A and five location-B trials, alternating repetitively (i.e., not randomized). The stimulation period lasted 30 s and the inter-trial period lasted 60 s. We collected up to two recording files from a given fly.

For the stimulation experiments in Fig. 5e-h, each fly experienced five left and five right LAL stimulation trials, presented in a random order. The stimulation period lasted 2 s and the inter-trial period lasted 30 s. We collected one recording file per fly.

### Data Acquisition

All timeseries data were digitized with a Digidata 1440A (Molecular Devices) at 10 kHz, except two-photon images, which were saved as tiff files using ScanImage at frequencies ranging from ∼4-10 Hz, as described above.

## Data Analysis

### Processing of behavioural data

The yaw, pitch, and roll angles of the ball were sampled at 50 Hz, and aligned to our imaging data files using the ball camera’s trigger signal. We shifted the acquired ball-position data backward in time by 30 ms due to our measured latency between the trigger pulse for acquiring a frame and when FicTrac finished processing the image.

For Extended Data Fig. 10b-c we used a 500-ms boxcar filter to smooth the forward walking velocity signal. For some analyses we excluded timepoints when the fly was standing still, which we defined as any moment when the fly’s filtered forward walking velocity was ≤ 1 mm/s. The fly’s virtual 2D trajectory was computed using the bar position, to estimate the fly’s heading, alongside the sideward and forward ball rotations to estimate the fly’s translational velocity. In Fig. 1, to visualize the relationship between neuronal phases and the fly’s orientation over time, we plotted the position of the bar on the arena (instead of the fly’s heading) since the EPG phase tracks the inverse of the fly’s heading (which is equivalent to the bar position)^4^. In Fig. 2, we flipped the heading direction *x*-axis to make it easier to compare with Fig. 1.

### Processing of menotaxis behavioural data

To analyze the fly’s menotaxis behaviour, we isolated straight segments (which we call *menotaxis bouts*) of the fly’s 2D virtual trajectory using the Ramer-Douglas-Peucker algorithim^56,57^ (Extended Data Fig. 2). This algorithm simplifies a set of *x*,*y* coordinates by iteratively reducing the number of points in the trace. The parameter *ε* determines the maximum allowed distance between the simplified and original trajectories. We then computed the fly’s displacement *L* for each segment of the simplified trajectory. For all analyses we used *ε* = 25 mm and only analyzed segments with *L* >200 mm. In other words, we analyzed menotaxis bouts where the fly displaced itself more than 200 mm (roughly equivalent to 70 body lengths), without deviating from its course by more than 25 mm (roughly 8 body lengths). Aside from bar-jump (i.e. virtual rotation) experiments––where we used the pre-jump heading angle as the fly’s goal angle––we defined the fly’s goal angle as the mean heading angle during each menotaxis bout. For this calculation, we excluded timepoints when the fly was standing still.

The values chosen for parameters *ε* and *L* were conservative, in that they tended to break up portions of the fly’s trajectory where one might have considered the fly’s goal to have remained unchanged into smaller bouts. We preferred this bias over the risk of potentially lumping two bouts together, where the fly’s true goal angles might have been different.

### Processing of imaging data

To correct for motion artefacts, we registered two-photon imaging frames using the CaImAn^58^ Python package. We defined ROIs for the left and right side of the LAL, the glomeruli of the bridge and columns of the fan-shaped body using a custom graphical user interface written in Python. ROIs were manually drawn using either the time averaged signal or the local correlation image of each *z* slice. In the case of the fan-shaped body, we used a semi-automated method to define columns as described previously^4^. Briefly, we first defined an ROI including the entire fan-shaped body. This ROI was then subdivided into 16 columns of equal angular size using two lines that defined the lateral edges of the fan-shaped body.

For each ROI, we defined ΔF/F_0_ as equal to (F - F_0_)/F_0_, where F is the mean pixel value of an ROI at a single time frame and F_0_ is the mean of the lowest 5% of F values. To align imaging data with behavioural data, we used a voltage signal of the *y* galvo flyback, which marks the end of an imaging frame, as an alignment point. For each imaging volume, the midpoint between the start of the volume’s first *z*-slice and the end of its last *z*-slice was used as its timestamp.

### Neuronal phase analysis

We computed the FC2 phase in the fan-shaped body using a population vector average^2,4^. We computed the EPG phase in the protocerebral bridge as described previously^4,38^. For each timepoint, we treated the glomeruli ΔF/F_0_ in the bridge as a vector of length 16 and took the Fourier transform of this vector. The phase of the Fourier spectrum at a period of 8.5 glomeruli was used as the EPG phase.

To overlay the FC2 or EPG phase with the bar position (Fig. 1, Extended Data Fig. 3), we subtracted from the phase its mean offset from the bar position. This offset was calculated, for each recording, by taking the mean circular difference between the phase angle and bar angle, excluding timepoints when the bar was in the 90° gap at the back of the arena or when the fly was standing still. In Fig. 1m-o and Extended Data Fig. 3d-f, we nulled the FC2 or EPG phase in the baseline period by subtracting its mean position, 1 s prior to the bar jump, from every sample point. In Fig. 1o and Extended Data Fig. 3f, we calculated the mean of this adjusted phase during the last 1 s of the open-loop period after a bar jump. To combine +90° and –90° bar jumps for analysis, the mean phase in the last 1 s of the open loop period was multiplied by −1 for −90° jumps.

In Fig. 1m-o, we imposed strict requirements for a bar jump trial to be included in the analysis. First, the bar jump needed to occur during a menotaxis bout (see *Processing of menotaxis behavioural data*). Second, we required that the fly return to its previous heading angle following a bar jump —i.e. trials when the mean bar position from 5 to 10 s after the start of the bar jump was within 30° from the mean bar position 5 s before the bar jump. Third, the bar needed to jump to a visible position on the arena. Finally, we only included trials when the FC2 or EPG population vector average amplitude was greater than 0.1. These criteria were sensible, in that they selected for trials where we could be confident that the fly’s goals had not drifted and that our neural signal estimates were of high quality. However, they were stringent enough that they led us to analyze only 7% of all trials. In Extended Data Fig. 3d-f, we eliminated the first two of these requirements, which lead us to analyze 54% of all trials.

To calculate the phase-nulled, population-averaged FC2 activity shown in Extended Data Fig. 3c, we followed a method described previously^38^,^4^. For each timepoint, we first interpolated the ΔF/F_0_ signal of the columns of the fan-shaped body to 1/10^th^ of a column using a cubic spline. We then circularly shifted the interpolated signal by the value of the FC2 phase of that timepoint. Finally, we averaged all the phase-aligned traces.

We used Scipy’s circmean function to compute the correlation between the EPG or FC2 phase and the bar position. For this calculation we excluded timepoints when the fly was standing still or when the bar was located in the 90º gap.

### FC2 stimulation analysis

To compare the effect of columnar stimulation of FC2 neurons across flies, we nulled the heading angle using the following procedure. For each fly, we computed its mean heading during a stimulation-A trial, excluding instances when the fly was standing still. We then took the mean heading across all stimulation-A trials and subtracted this value from the fly’s heading angle in all trials. The histograms in Fig. 2c-f used 10° bins and also excluded timepoints when the fly was standing still.

For Extended Data Fig. 5c, an ROI was considered inside the stimulation ROI if it had at least one pixel within the boundaries of the stimulation ROI scan path and was otherwise considered outside the stimulation ROI. In Extended Data Fig. 5d, we only analyzed ROIs that were outside the stimulation ROI. The change in the column ROI ΔF/F_0_ was computed by dividing the mean ΔF/F_0_ during the 30 s stimulation period and divided this number by the mean the ΔF/F_0_ during the 5 s before the stimulation. To calculate an ROI’s distance from the stimulation site (in number of ROIs), we first defined the stimulation site as the column ROI with the highest fraction of pixels inside the stimulation ROI. For each ROI we then computed its wrapped distance in number of ROIs. For instance, column ROI 2 and column ROI 15 have a distance of three (given that there are 16 columns in our analysis). Since our stimulation ROI could overlap with multiple column ROIs, in Extended Data Fig. 5d, there are no column ROIs with a distance of one.

In Extended Data Fig. 5e, to compute the stimulation location angle, we treated the fraction of pixels of each column ROI that were inside the stimulation ROI (see red colormap in Fig. 1c,d) as an array. Using this array, we computed the stimulation location angle with the same population vector average method used to compute the FC2 phase. We then took the mean difference between the two stimulation phases (A and B) for each fly. To compute the mean FC2 phase position during the stimulation period (Extended Data Fig. 5f-h), we excluded timepoints when the fly was standing still.

### LAL imaging analysis

To detect transient increases in LAL asymmetries (Fig. 5b-c), we first smoothed the right – left LAL ΔF/F_0_ signal using a Gaussian filter (σ = 200 ms). We then detected peaks in the filtered signal using the SciPy function signal.find_peaks. Peaks were defined as timepoints where the filtered signal was above 0.1 ΔF/F_0_ for at least 1 s, spaced from other peaks by at least 3 s, and had a prominence of one. To detected transient decreases in LAL asymmetries we flipped the right – left LAL ΔF/F_0_ signal and then applied the same algorithm. In Fig. 5c, we aligned the fly’s turning velocity and the right – left LAL ΔF/F_0_ signal to the timepoint of the peak neural signal and upsampled both the fly’s turning velocity and the right – left LAL ΔF/F_0_ to a common 100 Hz time base.

To plot the LAL activity as a function of the fly’ heading relative to its goal angle (Fig. 5d), we only analyzed data during menotaxis bouts (see *Processing of menotaxis behavioural data*). Because there is a ∼200 ms delay between a change in the fly’s heading and a change in the EPG phase^38^, we expected the LAL ΔF/F_0_ signal (which relies on this heading input) to be likewise delayed relative to behaviour. Therefore, in Figure 5d only, we shifted the LAL ΔF/F_0_ signal forward in time by ∼218 ms (2 imaging volumes) prior to relating the signal to the fly’s behaviour. We believe that this is the most appropriate signal to analyze, but our conclusions are the same if we do not apply this shift. For each fly, we calculated the mean LAL ΔF/F_0_ by binning the data based on the fly’s heading relative to its goal using 10° bins. We excluded timepoints when the fly was standing still.

### Processing of electrophysiological data

To detect spikes, we first filtered the membrane voltage (*Vm*) trace with a Butterworth bandpass filter. We then detected peaks in the filtered *Vm* trace above a specified threshold, spaced by > 5 ms, using the SciPy function signal.find_peaks. Although this criterion means we could not detect spike rates above 200 Hz, the activity levels of all our cells stayed well below this upper limit. Different cut-off frequencies and thresholds were hand selected for each cell so as to yield spike times that matched what one would expect from visual inspection of the data. To remove spikes from the *Vm* trace––for analyses of the membrane voltage in Fig. 3c-d, Extended Data Figs. 9d, 11c,d, 13b,c––we discarded *Vm* samples within 10 ms of a spike by converting those samples to NaNs.

When analyzing electrophysiological data in comparison to the fly’s heading or goal angle (Fig. 3c-f, Extended Data Figs. 9-12), we downsampled the cell’s *Vm* or spike rate to the ball camera frame rate (50 Hz) by either averaging the spike-rate or the spike-removed *Vm* in the time interval between two camera triggers. In Fig. 3b and Extended Data Fig. 10b,c, we plotted the spike rate using a 1 s boxcar average.

### Tuning curves

To generate the tuning curves in Fig. 3c-d, we binned the electrophysiological time-series data according to the fly’s heading, using 15° bins. We then calculated the mean spike rate and the spike-removed *V*_*m*_ for each bin. To estimate a cell’s preferred-heading angle, we fit the spike-removed *V*_*m*_ tuning curve with a cosine function, with the offset, amplitude and phase of the cosine (the phase is the resulting preferred angle) as fitting parameters. In performing this fit, we excluded timepoints when the bar was located in the 90° gap at the back of the arena because the EPG system is expected to track the fly’s heading less faithfully during these moments^2,38,42^. We used *<u>V</u>*_*<u>m</u>*_ rather than spike rate for estimating the cell’s preferred heading angle because *V*_*m*_ was much less modulated by the fly’s goal angle than the spike rate (Extended Data Fig. 11), and thus it was less likely to lead to goal-modulation-related biases in our estimate of the preferred heading angle.

For Fig. 3e-f and Extended Data Fig. 10-12, we only analyzed data from time points that contributed to a menotaxis bout (see *Processing of menotaxis behavioural data*). For each bout, we computed a relative goal angle by subtracting the cell’s preferred heading angle from the fly’s goal angle. Likewise, for each timepoint, we computed a relative heading by subtracting the cell’s preferred-heading angle from the fly’s current heading angle. We then calculated the mean firing rate (or spike-removed *V*_*m*_) binned by the fly’s relative goal angle using 45° bins (columns in Fig. 3f) and by the fly’s relative heading angle, also using 45° bins (*x*-axis in Fig. 3f). To generate tuning curves (except Extended Data Fig. 12), we excluded timepoints when the fly was standing still. In contrast, for Extended Data Fig. 12 we only included timepoints when the filtered forward walking velocity of the fly was below 0.5 mm/s (i.e. the fly was standing still) and the fly’s turning velocity was between –5°/s and 5°/s (i.e. the fly was not turning in place).

### Fitting the Tuning Curves

Because the data in Fig. 3f were binned according to relative heading and goal angles (relative to the preferred heading angle), which we denote here by *H*’ and *G*’, we expressed the PFL3 activity in the single-cell model as *f*(cos(*H*′) + *d*cos(*G*′ − *θ* + *φ*)), with *f*(*x*) = *a* log(1 + exp(b(x + c)). This form for *f*, which is called a softplus function, was suggested by examining the shifted spike-rate versus *V*_*m*_ curves in Extended Data Fig. 13c. We then fit the parameters (*θ* − *φ*), d, a, b and c by minimizing the squared difference between *f* and the data. The same value of *θ* − *φ* was used for each cell. The optimal parameters were *θ* − *φ* = −54°, *d* = 0.67, *a* = 22.09 Hz, *b* = 2.33 and *c* = −0.52. The connectomic analysis discussed in the next section indicates that the difference between the preferred heading and goal angles, *θ* − *φ*, is not actually the same for each PFL3 neuron, but this approximation was necessary because the recorded PFL3 neurons could not be identified as associated with specific glomeruli or compartments. A reasonable assumption is that the single fitted value for *θ* − *φ*, which was − 54°, would be approximately equal to the average value of this difference across the PFL3 population. However, this average difference is −66°(see the section on the full model). Several technical and biological reasons could account for the 12° difference between these values. For example, a misestimation the cell’s preferred-heading direction (see Tuning curves) could cause the measured *θ* − *φ* to be smaller than its average anatomical value.

### Spike-rate versus *Vm* curves

Extended Data Fig. 13b shows the relationship between the spike-rate and *V*_*m*_ (spikes removed) obtained from our whole-cell recordings. To generate this plot, we used the data shown in Extended Data Fig. 11 (i.e. we included timepoints when the fly was performing menotaxis and not standing still) but in this case, we binned the data according to the fly’s goal angle relative to the cell’s preferred-heading angle (using the same 45° bins) and according the cell’s *V*_*m*_ (4 mV bins). We used a cutoff of −46 mV, since at more depolarized membrane potentials spikes were not as well estimated. To include right PFL3 neurons in this analysis, we flipped the goal-heading-relative-to-the-cells’-preferred-heading values of right PFL3 cells prior to averaging across all cells.

To generate Extended Data Fig. 13c, in which the curves from Extended Data Fig. 13b are aligned, we shifted the curves for different goal directions along the horizontal (*V*_*m*_) axis by amounts determined to minimize the squared difference between the spike rates in each bin across the different goal directions. In other words, we computed the shifts that made the spike-rate curves for different goal directions maximally align. The resulting voltage shifts are plotted in Fig. 13d. The resulting aligned data was fit by a softmax function, *f*(*x*) = *a* log(1 + exp(β(*V*_*m*_ + γ)) (black curve in Extended Data Fig. 13c). The parameters β and γ of this fit are in different units than *b* and *c* for the fits in Fig. 3f, and it is the parameters of the latter fit that are used to build the full model, discussed next.

### Full PFL3 Model

For the full population model, the response of the right/left PFL3 cell innervating column *i* of the fan-shaped body (with *i* = 1, 2,…,12) was expressed as

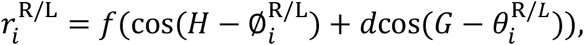

with *f* and the parameters *d, a, b* and *c* identical to what was described in the previous sections. The values of the preferred angles were obtained from the anatomy^15^ (Fig. 4b, Extended Data Fig.6d-e). For the preferred goal angles, we used the values *θ* = 15°, 45°, 75°, 105°, 135°, 165°, 195°, 225°, 255°, 285°, 315°, 345°. For the preferred-heading angles, we began by assigning angles to the 18 glomeruli across both sides of the protocerebral bridge, from left to right: 337.5°, 22.5°, 67.5°, 112.5°, 157.5°, 202.5°, 247.5°, 292.5°, 337.5°, 22.5°, 67.5°, 112.5°, 157.5°, 202.5°, 247.5°, 292.5°, 337.5°, 22.5°. These angles were projected down to the fan-shaped body using the wiring diagram shown in Fig. 4b. There are 18 bridge angles but only 16 of them are used for these projections because the left and right two outermost glomeruli (first and last two entries in the above list) are not innervated by PFL3 cells. For the PFL3 cells that innervate two glomeruli, we used the average of the angles corresponding to the two innervated glomeruli. The resulting preferred-heading angles are: *φ* = 67.5°, 112.5°, 157.5°, 157.5°, 202.5°, 225°, 247.5°, 292.5°, 337.5°, 337.5°, 22.5°, 67.5°, for the right PFL3 cells, and *φ* = 292.5°, 337.5°, 22.5°, 22.5°, 67.5°, 112.5°, 135.°, 157.5°, 202.5°, 202.5°, 247.5°, 292.5°, for the left PFL3 cells. These angles determine the directions of the vectors shown within the fan-shaped body compartments in Fig. 4b, with angles measured positive counterclockwise and the zero angle directly downward.

In the analysis described in the previous paragraph, we used glomerular angles implied by the Δ7 innervation of the protocerebral bridge (Extended Data Fig. 6c). Alternatively, we could have used glomerular angles based on the innervation of EPG neurons (Extended Data Fig. 6b). We opted to use the Δ7 scheme primarily because these angles predicted a population-level LAL signal that more closely fit our data. However, models using either sets of bridge angles produce qualitatively similar results. We also assumed that the PFL3 cells form twelve functional columns in the fan-shaped body due to anatomical considerations (Extended Data Fig. 7e-h). PFL3 neurons can, alternatively, be viewed as forming nine columns^14^. The model was also constructed assuming nine columns, and similar results were obtained as for the twelve-column model.

### Statistics

For Fig. 1o, to assess whether the FC2 phase changed relative to its initial position during a bar jump we performed a V-test^59^ (Rayleigh test for uniformity where the alternative hypothesis is a known mean angle, μ) with μ=0° (p=2.58×10^−3^). To assess whether the EPG phase tracks the bar during a bar jump we performed a V-test with μ=90° (p=2.62×10^−4^). The same tests applied to Extended Data Fig. 3 yielded μ=0° (8.95×10^−8^) for the FC2 phase and μ=90° (2.26×10^−4^) for the EPG phase.

For Fig. 2g, to assess whether the difference in flies’ mean heading direction for stimulation A and B was within the expected difference based on the stimulation locations in the fan-shaped body, we performed a V-test with μ equal the angular difference between the two stimulation location angles (Extended Data Fig. 5e). For flies expressing CsChrimson in FC2 neurons, this was μ=-173.4° (p=1.15×10^−3^). For control flies that did not express CsChrimson, this was μ=-164.8° (p=0.932). The expected difference of both groups is not exactly the same since the stimulation ROIs are defined manually without knowledge of the column ROIs (which are only defined later during the imaging analysis).

For Fig. 5h, to assess whether flies expressing CsChrimson in PFL3 neurons showed a change in ipsilateral turning velocity relative to control flies that only expressing jGCaMP7f, we performed a two-sided Welch’s *t*-test (p=1.93×10^−5^). To compare flies expressing CsChrimson in PFL1 neurons with control flies we used a two-sided Welch’s *t*-test (p=0.76).

## Notes

### Competing Interest Statement

The authors have declared no competing interest.

